# A Stochastic Modelling Framework for Cancer Patient Trajectories: Combining Tumour Growth, Metastasis, and Survival

**DOI:** 10.1101/2024.09.26.615161

**Authors:** Vincent Wieland, Jan Hasenauer

## Abstract

Cancer is a major burden of disease around the globe and one of the leading causes of premature death. The key to improve patient outcomes in modern clinical cancer research is to gain insights into dynamics underlying cancer evolution in order to facilitate the search for effective therapies. However, most cancer data analysis tools are designed for controlled trials and cannot leverage routine clinical data, which are available in far greater quantities. In addition, many cancer models focus on single disease processes in isolation, disregarding interaction This work proposes a unified stochastic modelling framework for cancer progression that combines (stochastic) processes for tumour growth, metastatic seeding, and patient survival to provide a comprehensive understanding of cancer progression. In addition, our models aim to use non-equidistantly sampled data collected in clinical routine to analyse the whole patient trajectory over the course of the disease. The model formulation features closed-form expressions of the likelihood functions for parameter inference from clinical data. The efficacy of our model approach is demonstrated through a simulation study involving four exemplary models, which utilise both analytic and numerical likelihoods. The results of the simulation studies demonstrate the accuracy and computational efficiency of the analytic likelihood formulations. We found that estimation can retrieve the correct model parameters and reveal the underlying data dynamics, and that this modelling framework is flexible in choosing the precise parameterisation. This work can serve as a foundation for the development of combined stochastic models for guiding personalized therapies in oncology.

## 1 Introduction

Millions of people are diagnosed with cancer each year (World Health Organization 2018), making it the second most common cause of death worldwide. For the successful invention of new prevention measures and improvement of treatments, it is crucial to deepen our understanding of the dynamics underpinning the progression of cancer on the molecular as well as the macroscopic level (Elmore et al 2021). Research solely based on clinical trials and traditional drug development pipelines faces the issue of high costs and ethical and regulatory roadblocks (Cui et al 2020). Consequently, it is of importance to leverage the large amount of longitudinal data that is routinely recorded over the time course of their disease. This ranges from the initial diagnosis to data collected during monitoring of the patients throughout the therapy and disease progression. Such data can include patient characteristics and pre-diagnostic diseases and interventions, bloodcount data, or histological and radiological reports. However, data collected in clinical routine is collected at random timepoints and often incomplete and unstructured. Therefore, it is of paramount importance to construct mathematical and computational methods that can use this data with the objective of facilitating the search for an efficacious cure personalized for each patient (Rahman et al 2022).

First groundbreaking attempts to aid this were made in the work of Collins et al (1956) and Schwartz (1961) concentrating on modelling the tumour growth as the main driving force of cancer. Soon, other mathematical models of differing complexity followed (Laird 1964; Steel 1977). Comprehensive reviews on the plethora of different tumour growth models can be found in (Gerlee 2013; Talkington and Durrett 2015; Tjørve and Tjørve 2017).

A realistic model of cancer progression does not only require the description of the size of the tumour, but also various other hallmarks (Hanahan and Weinberg 2000; Hanahan 2011) such as tumour vascularization, acquisition of mutations, and the spread of metastasis. Indeed, metastatic cancer is responsible for the majority of cancer-related deaths (Fares et al 2020). Due to further advances, in computing and cancer biology, larger computational models are possible, for example models for the mutation of key genes in cells (Gerlee 2013) or metastatic seeding (Franssen et al 2019; Nguyen Edalgo and Ford Versypt 2018).

Besides modelling the driving forces of cancer progression in the human body, tumour growth and metastatic seeding, it is important to monitor and evaluate the survival of cancer patients and model its dependence on different variables such as treatment decisions. Therefore, many countries and initiatives have established policies to raise cancer awareness and screening programs for early detection of cancers, leading to a large amount of time-to-event data for the history of cancer progression (Zhang et al 2023; Brito Fernandes et al 2024). Analysing this data with classical parametric survival models lacks the flexibility to capture the complex underlying dynamics of the cancer disease (Perera and Dwivedi 2020). Moreover, selecting an appropriate model can be challenging (Palmer et al 2023). In addition, more than one event or even the complete event history may be of interest, e.g. intermediate disease states may lead to different screening intervals. For these reasons multi-state models gained more attention in cancer screening evaluation and modelling cancer progression through different disease states (Cheung et al 2022; Uhry et al 2010). Still, they do not provide an understanding of the evolution of cancer progression in the patient.

As cancer modelling and screening capacities expanded, the speed of cancer therapy development increased; from classical chemotherapy evolving after Second World War, to drugs targeting specific molecules and lately to therapies using monoclonal antibodies and immune checkpoint inhibitors for the treatment of advanced or metastatic tumours (Arruebo et al 2011; Falzone et al 2018). Nowadays, the development personalized therapies that match the patient’s individual case is of main focus in the field of medical oncology (Falzone et al 2018). Also agent-based model gained attention in simulating the treatment decisions that need to be taken on the individual patient level (Mustapha et al 2016; Calvaresi et al 2020).

However, all of the afore-mentioned modeling approaches (Uhry et al 2010; Nguyen Edalgo and Ford Versypt 2018; Franssen et al 2019; Perera and Dwivedi 2020; Calvaresi et al 2020) mostly capture a single aspect of cancer patient trajectories. To efficiently guide the developments in modern oncology, mathematical models for the analysis of patient trajectories should not focus on tumour growth, metastatic seeding, or patient survival, but integrate all these processes to provide a holistic view on the disease progression in the patient. In addition, inter-individual differences of patients and the stochastic nature of the stochastic processes should be captured. To meet these requirements, mathematical models needs to integrate different types of governing equations to accommodate the nature of the individual processes.

Hence, research that probabilistically links tumour growth and metastatic seeding based on clinical routine data to develop a more holistic view on the patient gained attention (Heimann and Hellman 2000; Minn et al 2007; Gasparini and Humphreys 2022). One approach is the use of continuous growth based models. However, many of these are not flexible enough to be extended to adapt to different cancer entities or new processes (Isheden and Humphreys 2019). Another modelling approach tackling the task of providing a complete view on cancer progression is the use of discrete or continuous time multi-state Markov models. While Markov models can be easily extended, they quickly grow in complexity, which makes them computationally demanding and not suitable for practitioners (Uhry et al 2010; Isheden and Humphreys 2019). Joint models of longitudinal and time-to-event data simultaneously describe disease dynamics by a non-linear mixed-effect model and use a survival model for the patient health status, which depends on unobserved biomarker kinetics (Rizopoulos 2011; Wu et al 2012; Desmée et al 2017). While they provide a good framework for individual dynamic predictions based on patient’s covariates (Proust-Lima and Taylor 2009), they do not explicitly take into account the dependence of the different processes underlying the patient’s disease progression.

In this work, we propose a new type of combined stochastic model that addresses the challenge of being efficient and flexible while maintaining a holistic view. To fill the gap in high-level models that describe the patient’s trajectory based on routine clinical data in the existing model space, we provide a description of cancer progression and patient trajectories by expressing tumour growth, metastatic seeding, and patient survival with potentially stochastic processes. These three processes are then combined into a comprehensive model. The modelling framework is able to adapt to different assumptions about the underlying dynamics by exchanging the mathematical description of the underlying processes. In addition, the proposed modelling framework does not rely on a specific kind of data, but facilitates the utilization of the rich source of information about cancer patients collected in the clinical routine and keeps the flexibility to incorporate patient specific effects in each sub-process separately. To ensure efficiency in the inference of the model parameters from the data, likelihood functions are calculated analytically, as far as possible.

This paper is organised as follows. In Section 2 we introduce our combined model which involves the three main processes of cancer progression and their mathematical representation together with examples for precise model formulations. Furthermore, we validate our model by showing that we can mimick characteristics of real-world data sets. Next, we introduce the likelihood function of the model for parameter inference from a given data set in Section 3. In Section 4, we showcase the inference capabilities of the proposed model type in a simulation study with the example formulations from Section 2. First, the accuracy and efficiency of the analytically computed likelihoods are assessed in Section 4.2 and 4.3. We then show that our estimation procedure retrieves the correct model parameters in 4.4. Section 4.5 displays the flexibility of the modelling framework by using more complex representations for the tumour growth. Furthermore, we illustrate the ability of the model to discover the true underlying dynamics of the data through model selection (Section 4.6) and to identify treatment and covariate effects (Section 4.7). Finally, we conclude the manuscript with a discussion and an outlook on further extensions to apply the framework in different real-world data settings.

## 2 Modelling Framework

In order to provide mathematical models that can support and improve therapeutic decisions for patients, it is necessary to evaluate the progression of the cancer as well as the patient’s health trajectory, i.e. health status. For this purpose, we propose a modelling framework that considers three dynamic processes: (1) the growth of a primary tumour, (2) the seeding of metastasis, and (3) the patient survival. These processes are time-continuous but may have a discrete or continuous state space, rendering analysis and inference challenging. To ensure the feasibility of statistical inference, we will focus on models with tractable likelihoods.

### 2.1 Modelling of Cancer Growth, Metastatic Spread and Patient Survival

In the following, we will first describe models for each of the processes separately (Sections 2.1.1-2.1.2) and subsequently the combined model (Section 2.1.4).

#### 2.1.1 Modelling Tumour Growth

To model the growth of the primary tumour, we rely on continuous growth models for tumour progression, which gained increasing attention as flexible alternatives to standard multi-state Markov models (Yin et al 2019; Gasparini and Humphreys 2022; Strandberg et al 2023). These models express tumour size as a continuous function of time. Most commonly used are models for estimating the volume of the tumour, where tumour diameter measurements are recalculated into volume depending on assumptions of the spatial characteristics of the tumour (Talkington and Durrett 2015).

The first class of growth models we consider, consists of 1-dimensional ODEs modelling the total tumour size *S*(*t*) by

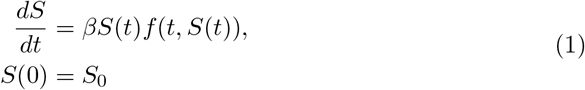

in which *β* denotes the growth rate and the function *f* gives the deviation from a standard exponential growth (Talkington and Durrett 2015). The choice of *f* might introduce additional model parameters. The term *S*(*t*)*f* (*t, S*(*t*)) then takes into account the underlying biological suppositions about the growth behaviour of the tumour., e.g. constant cell-division, growth only occurring on the surface, or saturation of the tumour environment. To ensure that a solution of the ODE exists, we assume the following (Teschl 2012).

##### Assumption 1

*The initial condition is positive, S*_0_ *>* 0, *and f* : ℝ → ℝ *is Lipschitz continuous on* [*S*_0_, ∞).

This general growth model (1) also includes the most commonly used formulations of tumour growth laws such as exponential growth, Gompertz growth, logistic growth, and power law representations (Gerlee 2013; Talkington and Durrett 2015; Spratt et al 1993). In this work, we will use exponential and Gompertz growth models.

##### Exponential Growth Model

The first model we use for modelling tumour growth is the simple exponential growth model, which resembles the assumption of a constant cell division rate independent of the tumour size. It was first applied to cancer in 1956 by Collins et al (1956).

By choosing *f* (*t, S*(*t*)) = 1 in the general growth model (1), the exponential growth of a tumour is represented by the following differential equation

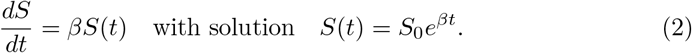

where *S*_0_ denotes the initial size of the tumour at detection *t*_0_ and is fixed across all patients. This model works well in the context of small tumour sizes and limited time, where no external effects lead to a halt and saturation in particular does not yet occur (Plevritis et al 2007; Abrahamsson and Humphreys 2016; Isheden and Humphreys 2019; Abrahamsson et al 2020; Gasparini and Humphreys 2022; Ocaña-Tienda et al 2024).

##### Gompertz Growth Model

As an alternative to the simple exponential growth model, we consider a Gompertz growth model. This model was introduced in Gompertz (1825) to analyse human mortality curves. As a model for biologic growth, it resembles the decrease of the growth of an organism due to factors like saturation (Wright 1926). Successfully applied to cancer by Laird in 1964 (Laird 1964), it became commonly used, especially for breast cancer (Steel and Lamerton 1966; Steel 1977; Norton 1988, 2005; Tabassum et al 2019).

We consider a Gompertz growth model with carrying capacity *K* expressed by the following ODE (Gerlee 2013).

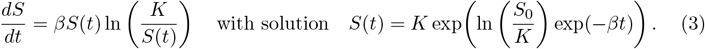

##### Gyllenberg-Webb Model

The modelling framework is not limited to 1-dimensional ODE models for tumour growth (1), but allows for more general continuous-time growth models. To demonstrate this, we consider in this work the physiologically structured tumour growth model by Gyllenberg and Webb (1990). This two-compartment model accounts for proliferating and quiescent cells, and is built on the assumptions that (1) actively proliferating cells can enter a quiescent state and (2) quiescence is more common in larger tumours. The model provides a generalization of Gompertz- and Bertalanffy-like S-shaped growth curves (Kozusko and Bajzer 2003; d’Onofrio et al 2011; Kuang et al 2018).

We consider the extended Gyllenberg-Webb model as proposed by (Alzahrani et al 2014), which accounts for proliferating cells *P*, quiescent cells *Q* and dead cells *R* and is described by

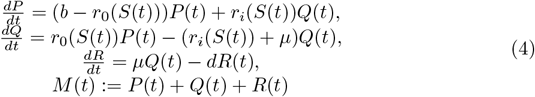

where cells proliferate at per capita rate *b >* 0 and die at rates *µ >*= 0. *M* gives the total tumour load. Additionally, cells transition between proliferating and quiescent compartment at rates *r*_*i*_(*S*), *r*_0_(*S*) subject to some general assumptions (Gyllenberg and Webb 1990; Kuang et al 2018). For this work we consider the example given in (Alzahrani and Kuang 2016)

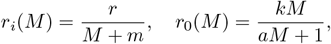

with *r* = 1, *k* = 2, *a* = 1, *m* = 2.

The Gyllenberg-Webb model is often used to include features related to resource limitation effects and yields a good starting point to explore treatment effect (Alzahrani et al 2014; Kuang et al 2018).

Since dead cells do not form metastases, we consider only proliferating and quiescent cells for the tumor volume *S*(*t*). However, it is also possible to base the metastasis process only on one part of the tumour, e.g. only on proliferating cells, if biologically justified for the tumour entity at hand.

Example growth curves for the three tumour growth models are depicted in Figure 1. A more exhaustive review on different forms of tumour growth laws with their respective advantages and interpretations can be found in (Marušić 1996; Gerlee 2013; Talkington and Durrett 2015; Kuang et al 2018), and the references therein.

**Fig. 1:**
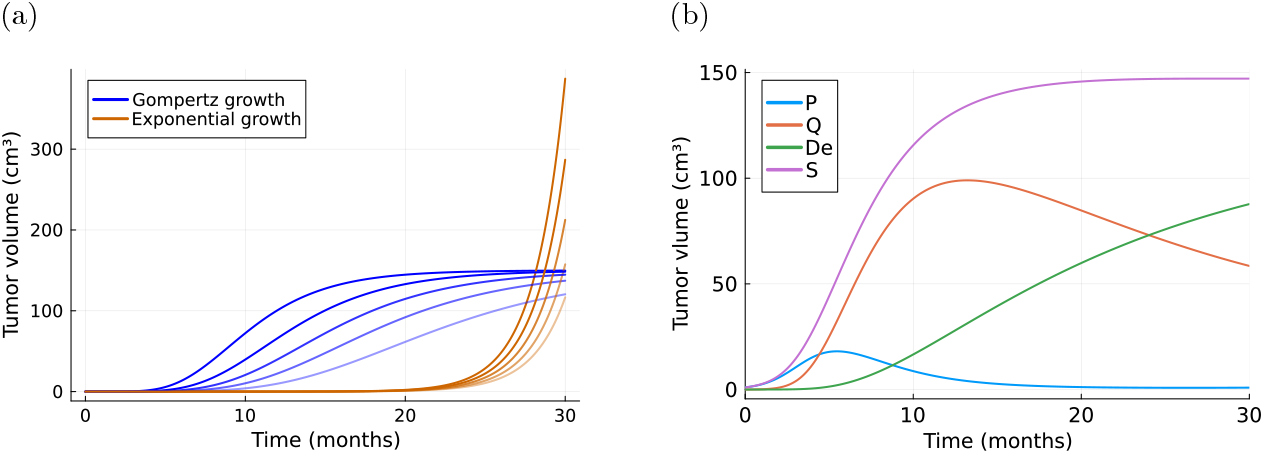
Simulations of tumour growth curves. 1a depicts example curves for exponential growth (2) and Gompertz growth (3). The initial tumour size is set to *S*_0_ = 0.065mm^3^ and saturation for the Gompertz growth to *K* = 150cm^3^. For the corresponding growth parameters we chose *β* ∈ {0.48, 0.49, 0.5, 0.51, 0.52} for exponential growth and *β* ∈ {0.14, 0.17, 0.2, 0.24, 0.3} for the Gompertz growth indicated by the shade of the lines. 1b shows an example curve for the Gyllenberg-Webb model (4), where we choose *P* (0) = 1, *Q*(0) = 0, *De*(0) = 0 and example parameter values [*b* = 1, *µ* = 0.05, *d* = 0.01, *r* = 1, *k* = 2, *a* = 1, *m* = 2].

#### 2.1.2 Modelling Metastatic Spread

Similar to the models of local spread to lymph nodes and metastatic seeding in the works of Gasparini and Humphreys (2022) and Isheden and Humphreys (2019), we exploit a stochastic model that models the number of metastases as 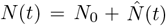. Here, 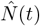 denotes an inhomogeneous Poisson point process. The rate of this Poisson process is chosen such that it accounts for the clinically established relationship between the size of the primary tumour and the spread of metastases (Koscielny et al 1984; Sopik and Narod 2018). More precisely, we choose an intensity function *λ*_*N*_ (*t, S*(*t*)) which depends on both time and tumour size and *N*_0_ denotes the number of metastases at detection time of the tumour. Thus the probability of *n* new metastasis seeding in a certain time-interval [*t*_*j−*1_, *t*_*j*_) is given by

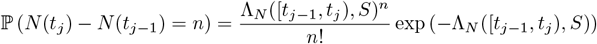

with

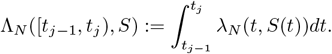

In order for this probability to be evaluable, we need the following assumption:

##### Assumption 2

*The intensity function is integrable, i*.*e. λ*_*N*_ (*t, S*(*t*)) ∈ *L*^1^(ℝ).

The rate of metastasis spread might depend on various factors, including number of cells, mutation stage, distance to vessel, and others. The intensity rate function *λ*_*N*_ can be formulated in a way that explicitly takes these dependencies into account. Here, we consider two specific example rates focusing on the relation between the rate of metastatic seeding and the size of the primary tumour.

##### Volume Based Metastatic Spread

Metastatic seeding is an exceedingly complex process involving many different steps (Arvelo et al 2016) and successful formation of metastasis depends on various not yet completely understood biochemical and genetic determinants that require research on their own (Hanahan 2011). As the number of cells which can leave the primary tumour to form metastasis depends on the size of the tumour, Bartoszyński et al (2001) proposed as a first model the intensity *λ*_*N*_ (*t, S*(*t*)) = *bS*(*t*). To account for the complexity of this process and other sources of metastatic seeding as well, we extend this and consider the following intensity

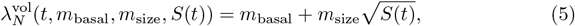

in which *m*_basal_ accounts for random metastatic seeding independent of tumour size and *m*_size_ is the coefficient of the effect of tumour size on the intensity. The dependence of metastatic seeding on observed characteristics can be incorporated by exchanging the constant parameters *m*_basal_, *m*_size_ by functions of various covariates such as age, medication, or others. The use of 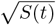 instead of *S*(*t*) showed more numerically robust behavior in our simulation study in Section 4. However, it can be interchanged by other functions as well.

##### Cell Division Based Metastatic Spread

One of the hallmarks of cancer is the capability of tumour cells to overcome hostile microenvironments when invading new systems (Hanahan 2011). Therefore, tumour cells need to be highly mutated to survive and establish a successful metastasis, which suggests a proportional relation of the rate of metastasis spread to the average number of mutations in the cancer cells and the rate of cancer cell division. Assuming a constant rate of mutation during cell division such a model has been proposed in Gasparini and Humphreys (2022) as an alternative to the volume based model intensity function (5). It is initially based on the model for lymph node spread by Isheden et al (2019), where they describe the volume of a spherical, dense tumour without cell death through the number of cell divisions *A*(*t*) and the size of a single cell *S*_cell_

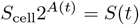

yielding for the number of cell divisions *A*(*t*) until a timepoint *t*

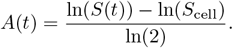

The rate of cell division in the tumour is then given by *A*^*′*^(*t*). Using this, the intensity is denoted by

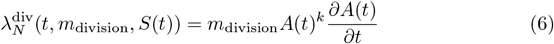

where the exponent *k* ≥ 1 being added for additional flexibility.

We compared the effect of the two tumour growth models on the metastasis process by simulating two datasets consisting of 500 patients. For both datasets we used the same parameters of the metastasis process and the death process and calculated the mean metastasis number. The visual difference depicted in Figure 2 showcases the influence of the choice of the tumour growth model on the process of metastatic seeding.

**Fig. 2:**
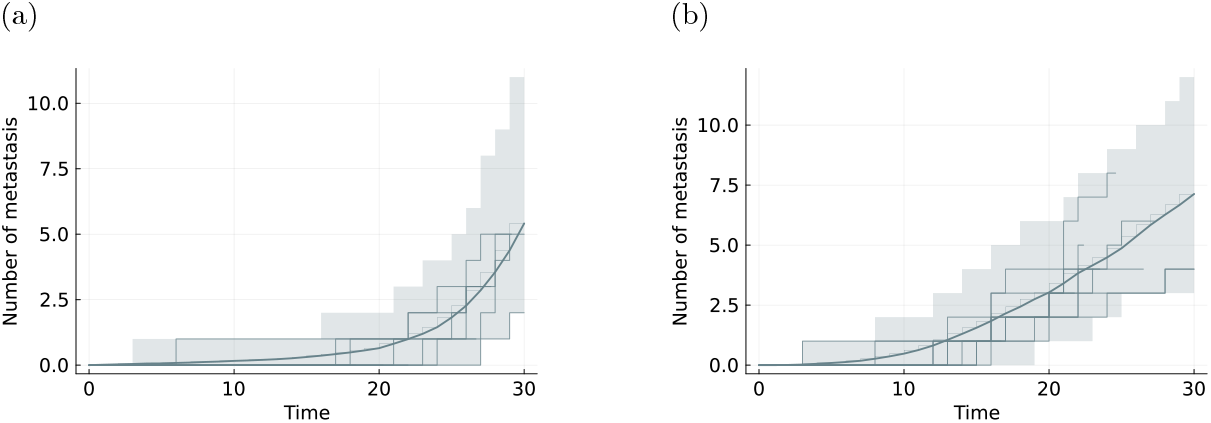
Simulations of the metastasis process. Visualization of the mean metastasis number (solid line) with 95% pointwise credibility intervals (shaded regions) and 10 realisations (stepped lines) from 500 patients using the volume based metastatic spread (5) with exponential tumour growth (2) in 2a and using the Gompertz growth model (3) in 2b. The rates *m*_basal_, *m*_size_ of the metastasis intensity are the same in both cases.

#### 2.1.3 Modelling Patient Survival

The third process which we consider is the survival of the patient. This process can be influenced by several confounding factors such as age or general health status and the underlying mechanisms still remain unclear (Boire et al 2024). However, the proximal causes of mortality in patients with cancer are often related to dysfunctional organs, for example, due to metastatic invasion (Fares et al 2020). A plausible assumption to resemble this relationship in our model is that the survival rate depends on the growth of the tumour and the number of metastases. A more thorough analysis would require model-based approaches. Additionally, cancer patients are observed over a limited time frame and most likely die due to the disease progression and not general aging. Therefore, we will leverage the simplifying assumption of having exponentially distributed survival times. This means that the time a patient will survive does not depend on the time the patient already survived since diagnosis. Formally, this is named the “memoryless property” and translates directly into the death process being a Markov process.

Hence, we model the survival of the patient by considering the death process *D*(*t*) to be an inhomogeneous Poisson point process that is stopped after the first jump occurs. The corresponding intensity rate is denoted by *λ*_*D*_(*t, N* (*t*), *S*(*t*)). Thus the probability to survive until some time *t >* 0 is given by

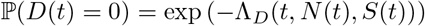

with

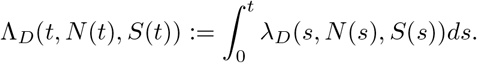

The intensity rate function *λ*_*D*_ depends on the tumour size and the metastatic load in the patient. As before we will assume integrability of *λ*_*D*_. We assume the following linear dependence on the tumour growth and metastasis number throughout the rest of this work

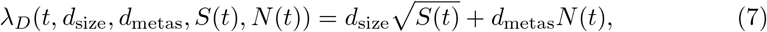

where *d*_size_ and *d*_metas_ quantify the influence of the corresponding process on the survival time.

#### 2.1.4 Combined Stochastic Model for Cancer Growth, Metastatic Spread and Associated Death

As the three previously processes are essential to describe and understand cancer patient survival, we combine them into one model providing a holistic description of the cancer patient. The resulting combined stochastic process, **X**(*t*) = (*S*(*t*), *N* (*t*), *D*(*t*))^*T*^, provides a novel formulation of a mathematical model for cancer progression. The Markovian nature of all three subprocesses ensures that the combined stochastic process still satisfies the Markov property. Furthermore, all subprocesses admit dependencies with each other, since the last observation timepoint is determined by the death process. A graphical exemplification using the example of breast cancer modeled with exponential tumor growth (2) and volume-based metastasis intensity (5) can be seen in Figure 3.

**Fig. 3:**
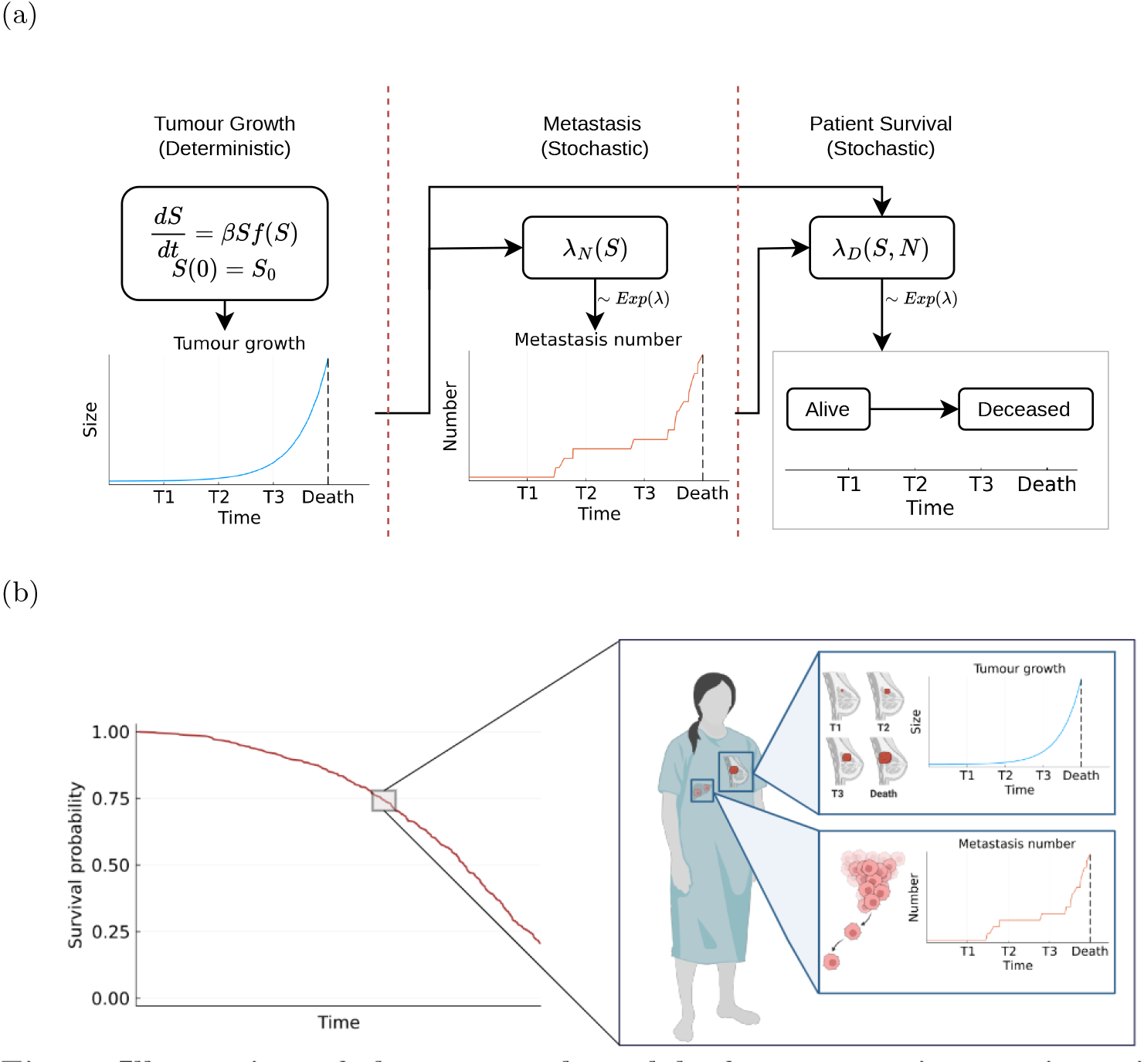
Illustration of the proposed model of cancer patient trajectories. 3a Outline of the three components of the model, the tumour size *S* with growth parameter *β*, the metastasis process *N* with rate *λ*_*N*_ (*S*) and the death process *D* with rate *λ*_*D*_(*S, n*) and their interconnections. 3b Simulation results for a population of 500 cancer patients using parameters reported in Supplementary Table S2: (left) Survival curve and (right) tumour growth and metastasis development for an exemplary patient.^1^

#### 2.1.5 Individual-Specific parameterisation

Cancer as a disease represents a complex ecosystem of hundreds of distinct types, which can vary substantially in their behavior. Even if one focuses on the analysis of one specific type of cancer, e.g. breast cancer, the progression varies between individual patients. In the above sections, we introduced rate parameters determining the dynamics of the process **X**(*t*) of interest, e.g. a tumour growth rate *β*. The combined stochastic model describes the stochastic evolution of cancer in a homogeneous population. Yet, it is well established that there are inter-individual differences which influence tumour growth rates *β*, metastasis development rates *m*_basal_, *m*_size_, *m*_div_ and other properties. Therefore, we allow the rates of the processes to depend on individual-specific, potential time-dependent covariates **Z**. The covariates for patient *l* are denoted by **Z**^*l*^(*t*), and might encode treatment, age, and weight.

##### Modelling Treatment Effect on Tumour Growth

Using the presented framework, we can investigate the effect of a treatment aimed at the growth of the primary tumour. We can model this by expressing the tumour growth rate *β* with a linear model of a basal growth rate and a covariate dependent effect. Both, the treatment status and other covariates that might influence the treatment effect are summarised in the covariate vector **Z**^*l*^(*t*) and the rate parameter is then given by

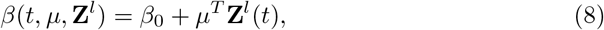

where *µ*^*T*^ is the vector of effects of treatment and covariates on the tumour growth. This linear relationship of covariates and model parameters can also be replaced by other functional representations.

With all model choices made, the parameters of interest are *β*_0_, *µ*^*T*^. We collect those together with the parameters of the other two processes, metastasis spread and patients’ survival, in the model parameter vector ***θ*** and denote the disease trajectory of patient *l* by **X**(*t*, ***θ*, Z**^*l*^).

### 2.2 Observation model

The state of the cancer progression within a patient cannot be observed continuously and without measurement noise. Accordingly, the model for the disease dynamics needs to be complemented by a model for the observation process. Here, we consider the case that the observation time points are determined by the visit time point of the patient to the hospital. Thus, the time points for the assessment of the size of the primary tumour and the number of metastases are independent from the two processes.

In clinical practice, the tumour size is mostly inferred by the use of imaging techniques rather than pathological measurements. Different factors such as radiographic imaging resolution or physician contouring preferences might lead to noise-corrupted data of the tumour size 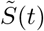 (Harshe et al 2023; Jakubowski et al 2012; Paquelet and Hendrick 2010). We represent this by adding a normally distributed random variable with size-dependent variance 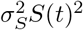 to the tumour size

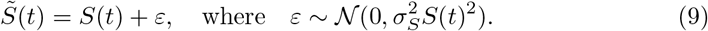

This assumption of normally distributed independent measurement noise can also be exchanged with different noise model assumptions. The application of such models needs to account for the measurement techniques and has to capture its characteristics in the observation function and the noise distribution. For the purpose of this work, the only requirement on the noise model is that it admits an analytical representation of the corresponding log-density function. To showcase the flexibility of our framework in the choice of the noise model, we provide an alternative formulation using log-normally distributed tumour size measurements in the Supplementary Information A.1.

Since the observation time points are uninformative and independent of the process, we assume that we observe the number of metastases 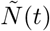 at each screening timepoint. In clinical practice, metastases, just like the primary tumour, must reach a certain detection threshold in order to be observed. However, under the assumption of constant growth rates for the metastases, this would only be a constant time delay in the detectability of events and will not influence the time passing between two events. Therefore, we assume that the metastasis process jumps as soon a metastasis reaches this threshold and we get observations 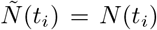 for the *i*-th observation time point.

As opposed to these observations at uninformative and process independent screening timepoints, the death is observed directly when it occurs. Hence, we observe the actual time of the transition of the death process from alive *D*(*t*) = 0 to dead *D*(*t*) = 1 and the last measurement timepoint always coincides with the timepoint of death *T*_*d*_ of the corresponding patient.

### 2.3 Implementation of the Combined Model

The code for the model implementation was written in the Julia programming language version 1.10.4 (Bezanson et al 2017). For the formulation and simulation of the differential equation for the tumour growth, we used the `Differentialequations.jl` package from the SciML ecosystem (Rackauckas and Nie 2017). On top of this, we implemented a version of the next reaction method based on Anderson (2007) to simulate the two inhomogeneous Poisson point processes depending on the solution of the differential equation.

### 2.4 Analysis of the Combined Model and Validation on Published Data

In the previous sections, we introduced a novel model for the description of clinical trajectories. To assess the plausibility of this model, we compare (1) the assumption about the subprocesses with real-world data on patient trajectories, and (2) the properties of the overall model with real-world data from clinical trials.

#### Assessment of assumptions on subprocess’ properties

So far we introduced a modelling framework and presented simple models for the subprocesses. While these subprocess models can be easily replaced, we hypothesize that some assumptions might appear more restrictive than they actually are as the coupling within the combined model allows for additional degrees of freedom.

To assess this hypothesis, we consider the assumption of a simple Markov processes for survival. On first glance, this might suggest that the survival tile follows an exponential distributions, which has been shown to be unrealistic for many tumor types (Baghestani et al 2016; Klakattawi 2022). Yet, as the simple survival process is coupled with a growth model for the tumour size and a metastasis process, the distribution of the survival times does not admit a simple exponential distribution as seen in the histograms of the overall survival times (Supplementary Figure S1). Since for each patient the overall survival time consists of a sequence of inter-transition times with survival rates being influenced by the tumour growth and the jump process of metastasis count, one faces a sequence of exponentially distributed times with differently time-varying rates. To investigate the distribution of the overall survival we fitted a general survival time distribution using a maximum-likelihood approach for censored data. Namely, we used the generalized Gamma distribution^2^ to describe the survival times obtained by simulating two models since it inherits common survival time distributions such as the exponential distribution, the Weibull distribution and the Gamma distribution as special cases (Box-Steffensmeier and Jones 2004). The first model we used combined exponential growth with the proportional metastasis intensity and the second used Gompertz growth to describe the tumour size. The survival time distribution for the first model closely resembles the a generalized Gamma distribution (Supplementary Figure S2). The survival time distribution for the second model showed to be close to a Weibull distribution based on the one-sample Kolmogorov-Smirnov test (K-S test) and Pearsons chi-squared test (Agresti 2013). The Weibull distribution is commonly used in describing survival of cancer patients and treatment effects on it (Baghestani et al 2016; Plana et al 2022; Klakattawi 2022).

To study the capabilities of the model to describe real-world observational data, we consider the ACT2-trial (Hagman et al 2016). In this trial, colorectal cancer patients with KRAS mutations were treated with ACT2. The dataset has been published Cancertrials.io (2022) and was previously fitted with a Weibull distribution Plana et al (2022). The Gompertz growth based model was able to mimick the survival curve of this trial as shown in Figure 4. MLE-based distribution fitting showed that a Gompertz distribution with parameters *k* = 2.7, *λ* = 26.7 describes the survival data from the ACT2-trial and the survival times obtained by our model with the parameters given in 4b based on the K-S and Pearson’s chi-squared test.

**Fig. 4:**
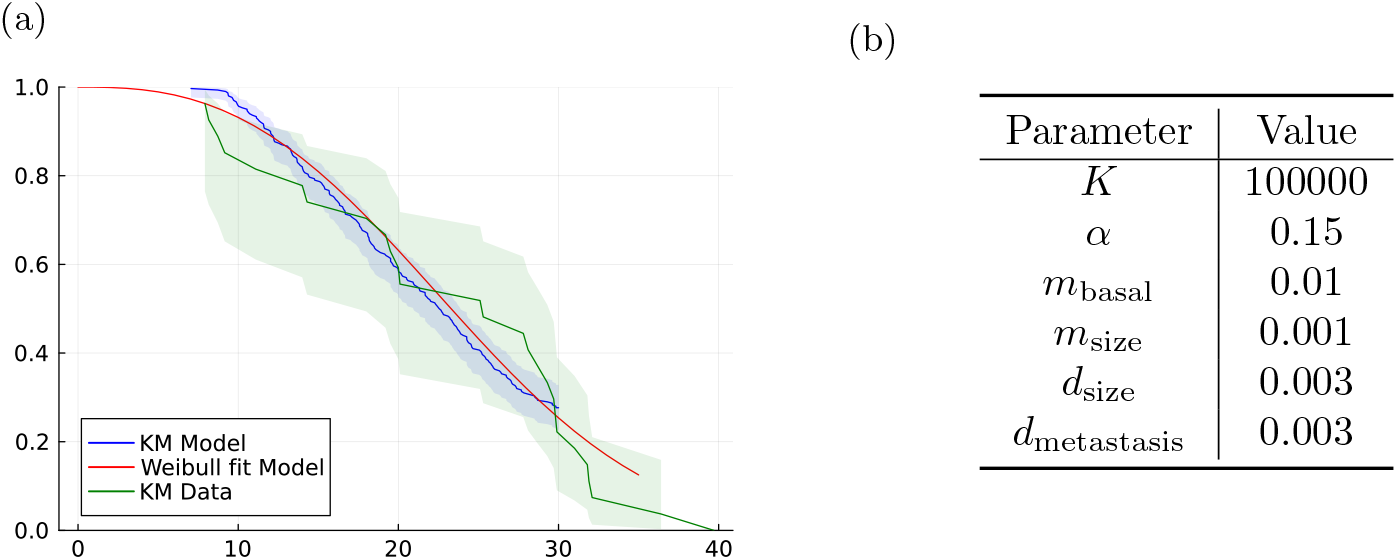
Assessment of survival times distribution. 4a provides a visualization of the estimated survival curve of model (M4) combining Gompertz growth with exponential proportional metastasis intensity (blue) against the estimated survival curve of 33 patients from the ACT2-trial (green). Additionally, the survival curve of a Weibull distribution with parameters *k* = 2.7, *λ* = 26.7 (red) is plotted. 4b provides the parameter values used to simulate the data from the model.

#### Assessment of proposed model structure

To validate the proposed model structure in the context of real-world data, we investigate whether the model can recover characteristics of real-world datasets. For this purpose, we considered the work of Engel et al (2003), in which they analyzed 12423 patients with breast cancer from the Munich Cancer Registry (MCR)^3^ from 1978 to 1996. They stratified the patients into the four T stages pT1, pT2, pT3, pT4 of the TNM classification system depending on the initial size of the primary tumour at time of diagnosis. As one of the results they reported that the overall survival following diagnosis varies significantly across these groups (Figure 1 in (Engel et al 2003)). Whereas survival after metastasisation appears to be independent of the pT category, indicating an almost homogeneous growth of metastases (Figure 3 in (Engel et al 2003)). The model combining exponential tumour growth with the cell division based metastasis rate (6) can reproduce these findings. We simulated 4 datasets of 1000 patients which only differed in the initial tumour size but not the model parameters. Initial tumour sizes were chosen to represent the four pT categories with corresponding diameters 10mm, 35mm, 75mm and 120mm and the model parameters are given in Supplementary Table S1. Only using different initial tumour sizes the model is capable of representing the finding that survival after metastasisation is nearly independent of the primary tumour size at time of diagnosis, but over survival differs significantly across the four groups. Additionally, we used the WebPlotDigitizer software (Ankit 2025) to recreate the survival curves for overall survival and survival after metastasisation from Figure 1 and Figure 3 in (Engel et al 2003). Figure 5 shows that the survival curves obtained from the simulated datasets with our proposed model show a consistent behavior with the ones reported.

**Fig. 5:**
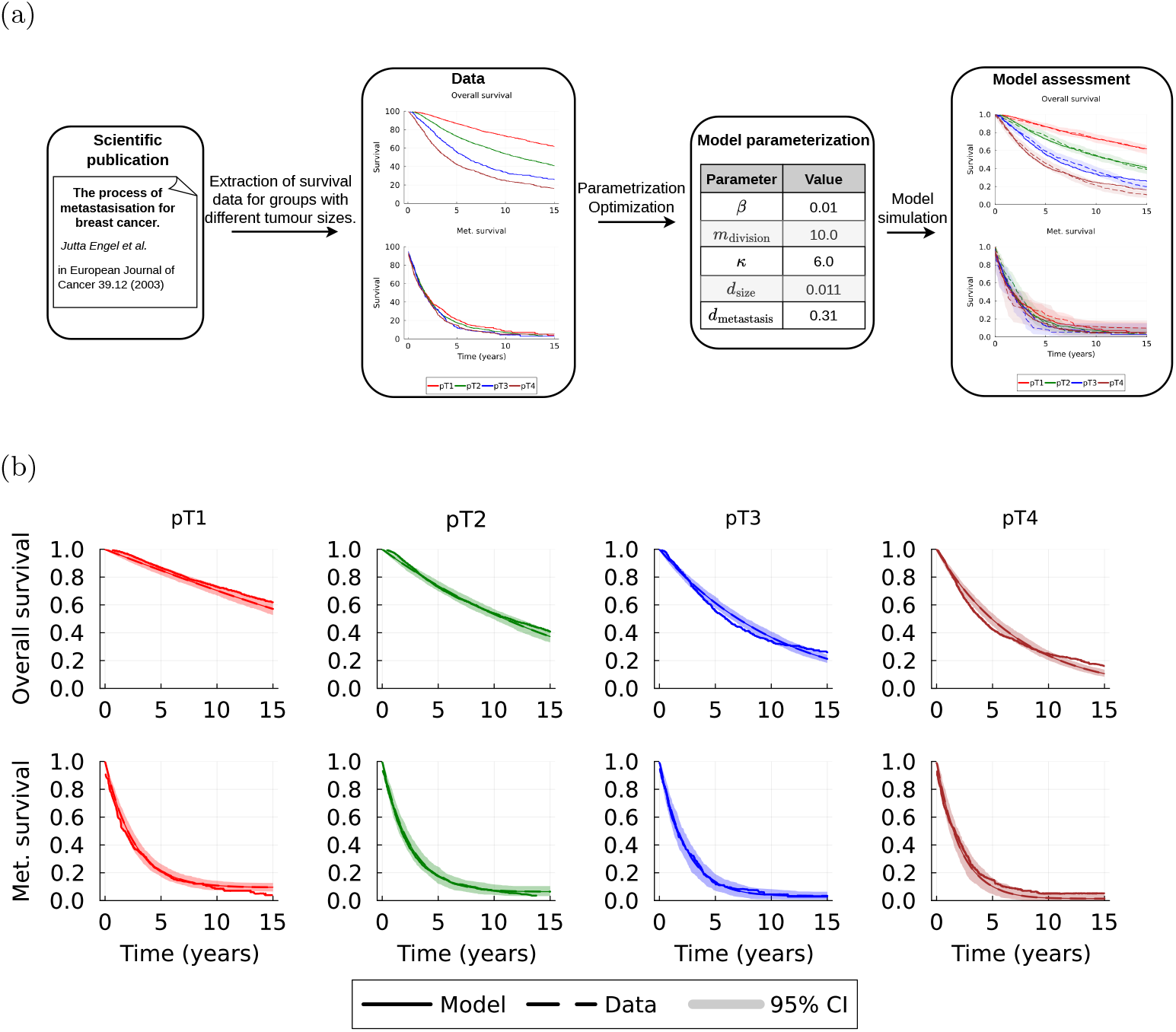
Assessment of proposed model structure. (a) Visual outline of model assessment performed using the dataset by Engel et al (2003) on overall survival and survival after metastasisation of breast cancer patients across different tumour sizes at time of diagnosis. (b) Comparison of observed (Data) and simulated (Model) survival curves. The simulation results are indicated using the mean survival curve (line) and the 95-percentile interval (shaded area) from 200 simulations of our model.

## 3 Parameter Inference

The combined stochastic model for a cancer patient depends on several model parameters determining the tumour growth rate, metastatic seeding rate, and survival rate. To correctly assess disease progression, impact of treatment effects and related properties, it is necessary to infer these parameters characterising the evolution of a patient from their corresponding clinical data. This can be achieved using likelihood-based and likelihood-free approaches. Likelihood-free such as approximate Bayesian computation are computationally expensive (Schälte et al 2022), leaving likelihood-based approaches as the method of choice. Yet, they require a likelihood formulation which admits computationally efficient evaluation.

Here, we derive such formulations for the aforedescribed combined stochastic model: a fully analytical expression and an expression which admits efficient numerical integration. Subsequently, we describe how these formulations are used for parameter optimisation and uncertainty analysis.

### 3.1 Computation of the Likelihood Function

We consider the inference of the model parameters from the clinical data for a population of *K* patients, 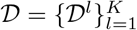. As the clinical datasets for individual patients are independent the likelihood can be factorized as follows

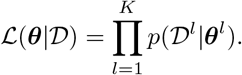

In the following, we outline without loss of generality how the likelihood contribution from patient *l, p*(*𝒟* ^*l*^ |***θ***^*l*^, **Z**^*l*^), can be evaluated. For ease of notation we omit the patient index in the rest of the section as well as the possible dependence on patient specific covariates available in the data.

For a single patient, the likelihood describes the probability of observing the corresponding dataset 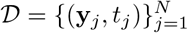, consisting of observations

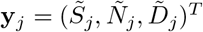 at timepoints 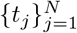, given a parameter vector ***θ***,

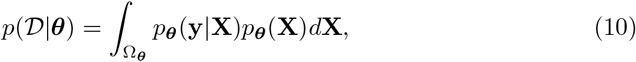

where Ω_***θ***_ is the full state-space of the combined stochastic process **X** for a given parameter vector ***θ***. The first term in the integral is the likelihood of the observation model and the second term the likelihood contribution of the hidden process.

Due to the Markov property of **X**, the observation timepoints divide the sample path of the process into mutually independent parts. Therefore, we obtain the likelihood for one patient by factorising over the observation timepoints.

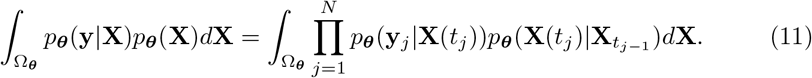

#### Likelihood of the Observation Model

The first term in the product on the right-hand side of (11), *p*_***θ***_(**y**_*j*_|**X**(*t*_*j*_)) = *p*(**y**_*j*_ |**X**(*t*_*j*_), ***θ***), is the probability of observing **y**_*j*_ given the actual state **X**(*t*_*j*_) = (*S*(*t*_*j*_), *N* (*t*_*j*_), *D*(*t*_*j*_))^*T*^ of the process at observation timepoint *t*_*j*_. Due to the assumptions made in Section 2.2, we exactly observe the number of current metastases and the death timepoint *T*_*d*_ of the patient. The latter is equivalent to observing the current survival status of the patient *D*(*t*_*j*_). This yields 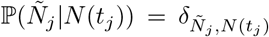 and 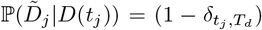, where *δ*_*i,j*_ denotes the Kronecker delta. Hence, the only contribution to the likelihood of the observation process is given by the measurement noise in the tumour size measurements. Therefore, we can write the likelihood contribution of the observation model as

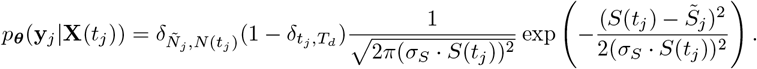

This yields that the integral in (11) vanishes on every possible path of **X** except for the set of paths exactly matching the observations of metastasis process and survival which we denote by 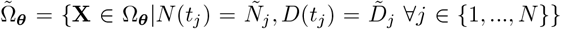. Since we assume that the deterministic growth process for the primary tumour is uniquely defined by the corresponding growth rate parameters in ***θ*** the observation likelihood is independent from the realized path and we can then rewrite

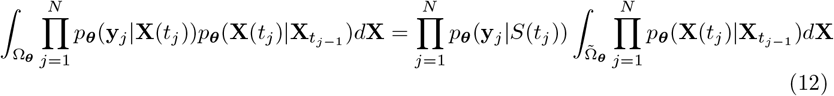

Given the initial condition *N*_0_, *D*_0_ the second part of the likelihood (12), the process likelihood, is given by

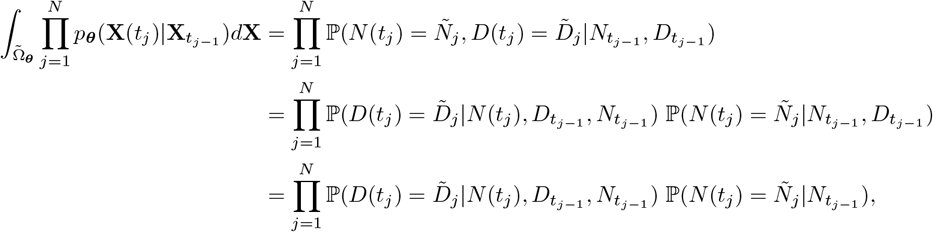

which decomposes the likelihood of the process dynamics into a likelihood contribution of the metastasis counting process and one of the death process. Both are investigated in more detail in the next steps. Note that on the right hand side and in the rest of the section we omit the dependence on ***θ*** for notational brevity.

#### Likelihood of the Observed Number of Metastasis

Since the metastasis number is not bounded from above and we are assuming an underlying time-inhomogeneous Poisson counting process, the likelihood contribution of the metastasis process is given by

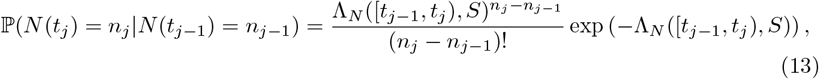

where *S* is the deterministic tumour growth process, i.e. the solution of (1) uniquely defined by the model parameters and the initial condition corresponding to the tumour growth.

#### Likelihood of the Observed Timepoint of Death

For the likelihood contribution of the death process, we need to marginalise out the unseen timepoints at which new metastases occur between two observation timepoints *t*_*j−*1_, *t*_*j*_, leading to the following results.

##### Theorem 1

**(Survival likelihood)** *Let S*(*t*) *be the solution to the ODE* (1) *satisfying Assumption 1; let λ*_*N*_ (*t, S*(*t*)) *be the intensity function of a Poisson point process N* (*t*) *admitting Assumption 2; let λ*_*D*_(*t, S*(*t*), *N* (*t*)) *be the intensity function of D*(*t*) *given by* (7). *Then the probability of survival given the occurrence of m new jumps of N* (*t*) *in a time interval* [*t*_*j−*1_, *t*_*j*_) *is given by*

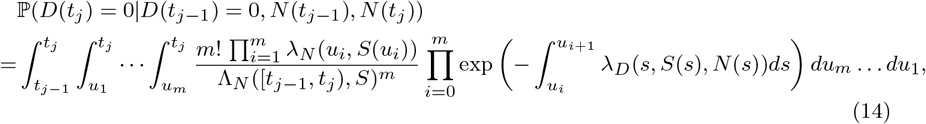

*where u*_0_ := *t*_*j−*1_ *and u*_*m*+1_ := *t*_*j*_.

Using this, one can directly deduce the probability of death occurring at time *t*_*j*_ given that *m* new metastasis occurred after the last observation at *t*_*j−*1_.

##### Corollary 2

**(Death likelihood)** *Under the same assumptions as before, we get*

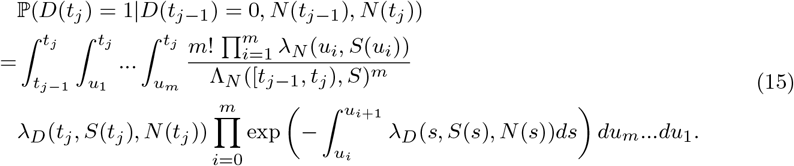

The proofs of these two results can be found in the Supplementary Information D.

##### Analytical Likelihood Formulation for Models with an Exponential Tumour Growth Law

For models based on an exponential tumour growth law as described in Section 2.1.1 combined with any of the two metastasis processes and the death process presented in Section 2, we were able to solve the nested integrals involved in equations (14) and (15) analytically. This yields a completely analytically available likelihood function for these models. The precise computation and simplification of the analytical solutions were done using symbolic computation in the Wolfram Mathematica computer algebra system (Wolfram Research, Inc. 2024).

For the model choices for which an analytical solution was not achievable, we leveraged efficient numerical integration schemes to solve the nested integrals and obtain a numerical approximation to the likelihood function that could then be further used for the optimisation (Leader 2022).

##### Likelihood calculation in the case of missing data

Models designed for the use with clinical data face several challenges, notably the use of irregularly sampled, noisy and partially missing measurements. The first is addressed by the use of different observation noise models as pointed out in Section 2.2. Corresponding to the data at hand such noise models can be applied to each of the subprocesses of the combined model. Irregularly sampled data are naturally supported by the model as it does not assume any structure on the distribution of measurement times and the likelihood for the proposed modelling framework can be factorized over timepoints, see Eq. (11), this is inherently addressed and one of the advantages of a Markovian model. Partially missing timepoints, where only measurements of some of the subprocesses are provided, can be naturally addressed as well, by marginalising over the missing components of the model. Let the measurement of the *i*-th component of the process *X* be missing at timepoint *t*_*j*_, then the likelihood reads as

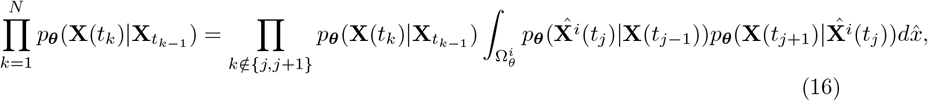

where 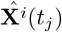 denotes the process with the *i*-th component being 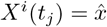.

Given that, if a measurement timepoint exists, one knows at least the survival status of the patient we consider the case of missing tumour size measurement and missing metastasis number measurement. In the first case, marginalising over all possible tumour sizes will just yield an likelihood value of the observation model equal to 1. Hence, we can just omit the likelihood of the observation model at the timepoint *t*_*j*_. In the case of a missing metastasis number measurement, the integral in (16) will become a sum.

If the patient dies at timepoint *t*_*j*_, i.e. 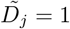, we can write

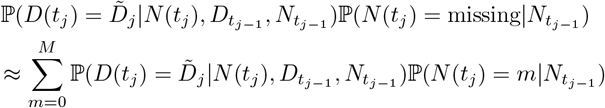

where the cut-off *M* is chosen to be the highest number of metastasis we observed in the data.

If the patient does not die at *t*_*j*_ the marginalised likelihood will be

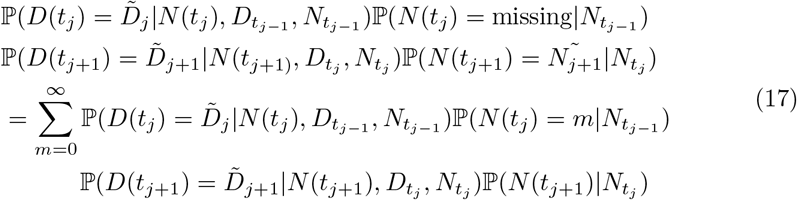

where all summands with *m* ∉ [Ñ_*j*−1_, Ñ_*j*_] are zero.

For the case where the patient does not die in the interval [*t*_*j−*1_, *t*_*j*+1_] Eq. (17) simplifies even more. Leveraging the Chapman-Kolmogorov equation (Pavliotis 2014) one can just skip the process likelihood at timepoint *t*_*j*_ and at the next timepoint compute *p*_***θ***_(**X**(*t*_*j*+1_|**X**(*t*_*j−*1_).

### 3.2 Parameter optimisation

The maximum likelihood estimate ***θ***^MLE^ of the model parameters is obtained by maximising the likelihood of observing the data given the model, computed in the sections above. In this work, we determine the maximum likelihood estimate by minimising the negative log-likelihood yielding

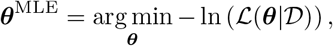

where ***θ*** denotes the vector of model parameters characterizing the combined stochastic process **X**. The minimization is done with computational optimisation techniques that need to be chosen according to the likelihood formulation at hand. For the case of analytical likelihood functions, we additionally compute gradients to leverage a gradient-based minimisation algorithm such as gradient descent or quasi-Newton methods, whereas with the numerical approximation to the likelihood we fall back to non-gradient based methods such as simulated annealing (Press 2007).

### 3.3 Uncertainty Analysis

For models which provide an interpretation of the model parameters, an assessment of how certain we are in the parameter estimates is meaningful. We assessed this parameter uncertainty by approximating the posterior distributions of the parameters *p*(***θ*** |D). This was done using a Markov chain Monte Carlo (MCMC) sampling algorithm (Andrieu et al 2003), to approximate the posterior distribution by a large number of samples and compute corresponding credibility intervals for ***θ***^MLE^. More precisely, we used an adaptive parallel tempering algorithm (Vousden et al 2015) to run several independent Markov chains and discarded the first half of the chain as a burn-in.

### 3.4 Implementation of the Inference Pipeline

As the code for the model simulation, we also implemented the likelihood-based inference procedure purely in the Julia programming language. Analytical solutions of the likelihoods were written down precisely and gradients were, if possible, computed by automatic differentiation (Revels et al 2016). For the numerical approximations of the likelihood we applied the numerical integration schemes from the Cubature.jl package (Johnson 2020). In particular, we leveraged an p-adaptive numerical integration scheme based on Clenshaw-Curtis quadrature rules Clenshaw and Curtis (1960). This can be seen as an expansion of the integrand in terms of Chebyshev polynomials and is well suited for accurately calculating low-dimensional integrals (Johnson 2010). We chose a relative error tolerance of 1*e*^*−*8^. While in principle differentiable with respect to the model parameters, the implementation using Cubature.jl for a robust and fast evaluation of the nested integrals does not allow for differentation.

Following comprehensive testing, we decided for running multiple starts of a local optimisation algorithm. For gradient-based optimisation in the cases where the gradient was available, we employed the limited-memory Broyden-Fletcher-GoldfarbShanno (LBFGS) algorithm Liu and Nocedal (1989). For gradient-free optimisation, we used the simulated annealing with box constraints (SAMIN) algorithm (Kirkpatrick et al 1983; Goffe 1996). Both are implemented in the Optim.jl package (Mogensen and Riseth 2018). For improved numerical stability, model parameters were transformed to log-scale during the optimisation.

For Markov chain Monte Carlo sampling, we used an adaptive parallel tempering algorithm (Vousden et al 2015) as implemented in the Python package pyPESTO (Schälte et al 2023) that was interfaced from Julia.

All computations were conducted on 10 cores of an AMD EPYC 7F72 3.2 GHz processor with 20GB of RAM. In order to ensure reusability and reproducibility, we made the code for model simulations and experiments together with the artificial data and results generated for this paper available at Zenodo (https://doi.org/10.5281/zenodo.13839104).

## 4 Evaluation of Parameter Inference

In this manuscript, we introduced a model class for the joint analysis of tumour growth, metastatic development, and patient survival. In the following, we assess the computational complexity of likelihood evaluation and parameter estimation. Furthermore, we evaluate the agreement of numerically and analytically computed likelihood values, the accuracy of parameter inference for different model structures, and the distinguishability of different parameterisations and model structures. Overall, we consider 5 different example models consisting of different combinations of growth and metastasis processes as summarized in Table 1. The death process is the same for each of the models.

**Table 1:** Model formulations. Overview of the models used for the simulation study in Section 4.

| Model name | Tumour growth process $S(t)$ | Metastasis process intensity function |
| --- | --- | --- |
| (M1) Exponential proportional model | $S(t) = S_0 \exp(\beta t)$ | $\lambda_N^{\text{prop}}(t) = m_{\text{basal}} + m_{\text{size}} \sqrt{S(t)}$ |
| (M2) Cell division model | $S(t) = S_0 \exp(\beta t)$ | $\lambda_N^{\text{div}}(t) = m_{\text{division}} \beta \left( \frac{\beta t}{\ln(2)} \right)$ |
| (M3) Gompertz model | $S(t) = K \exp \left( \ln \left( \frac{S_0}{K} \right) \exp(-\beta t) \right)$ | $\lambda_N^{\text{prop}}(t) = m_{\text{basal}} + m_{\text{size}} \sqrt{S(t)}$ |
| (M4) Gyllenberg-Webb model | $S(t) = Q(t) + P(t) + De(t)$ | $\lambda_N^{\text{prop}}(t) = m_{\text{basal}} + m_{\text{size}} \sqrt{S(t)}$ |
| (M5) Treatment effect model | $S(t) = S_0 \exp(\beta \mathbf{Z}(t)t)$ | $\lambda_N^{\text{prop}}(t) = m_{\text{basal}} + m_{\text{size}} \sqrt{S(t)}$ |

### 4.1 Study Design

For each of the models, we created a dataset of 500 patients under the following experimental conditions. Using the corresponding model description, we simulated the whole trajectory until the death of the patient or a maximum time of 30 months and read out observations at timepoints *t*_0_, …, *t*_*n*_, corresponding to monthly visits of the patient. For the simulation of the tumour growth given by (1) we assumed that tumours originate from a single spherical cell of volume *S*_cell_ corresponding to a diameter of *d*_cell_ = 0.01mm, but are detectable only once they have reached a volume *S*_detection_ corresponding to a diameter of *d*_detection_ = 0.5mm (Gasparini and Humphreys 2022; Hao et al 2018). Hence, the first observation timepoint *t*_0_ of a patient corresponds to the time between offset of the tumour and diagnosis. However, under the assumptions that the initial volume and detection threshold are known and given the model parameters, we could calculate the time since the offset of the tumour *t*_0_ for each patient, based on the first observation. Therefore, without loss of generality we assumed that tumours are detected as soon as they reach the detection threshold, i.e. *S*_0_ = *S*_detection_ = 0.065mm^3^ for all patients and omitted the constant linear time shift and set *t*_0_ = 0 for all patients. Additionally, the Markov property of the metastasis process ensures that the gain of new metastasis is independent of how many metastasis the patient has developed so far. Therefore, without loss of generality we initialized the patient simulations with zero metastasis at the first observation timepoint, i.e. *N*_0_ = 0.

### 4.2 Numerical and Analytical Likelihood Values Agree for Exponential Growth Based Models

In Section 3.1, we formulated the likelihood function and described the availability of analytical solutions for certain model formulations. Here, we examine the solution of the likelihood function for model (M1) and (M2), which are based on the exponential tumour growth law. The combination of the exponential tumour growth law with the volume-based metastasis intensity enables the computation of the accumulated metastasis intensity rate as

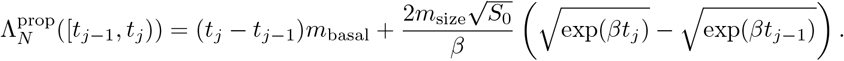

For the second model the use of the cell division based intensity rate for the metastatic spread (6), yields

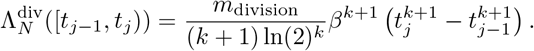

These accumulated metastasis intensities directly provide analytical expressions for the metastasis likelihood (13). Moreover, both model formulations enable the computation of analytical solutions for the nested integrals in the likelihood contribution of the death process equation (14) and equation (15). The precise expressions for the number of new metastasis *m* ∈ {1, 2, 3, 4, 5} are given in the Supplementary Information E, where the upper bound on *m* was chosen due to the maximal number of new metastasis observed in one inter-observation time interval in the simulated datasets. Artificial data was simulated using parameters provided in Supplementary Table S2

For both models evaluating the resulting analytical negative log-likelihood function shows a complete agreement with the corresponding approximated negative log-likelihood function obtained by numerical integration as seen in Figure 6.

**Fig. 6:**
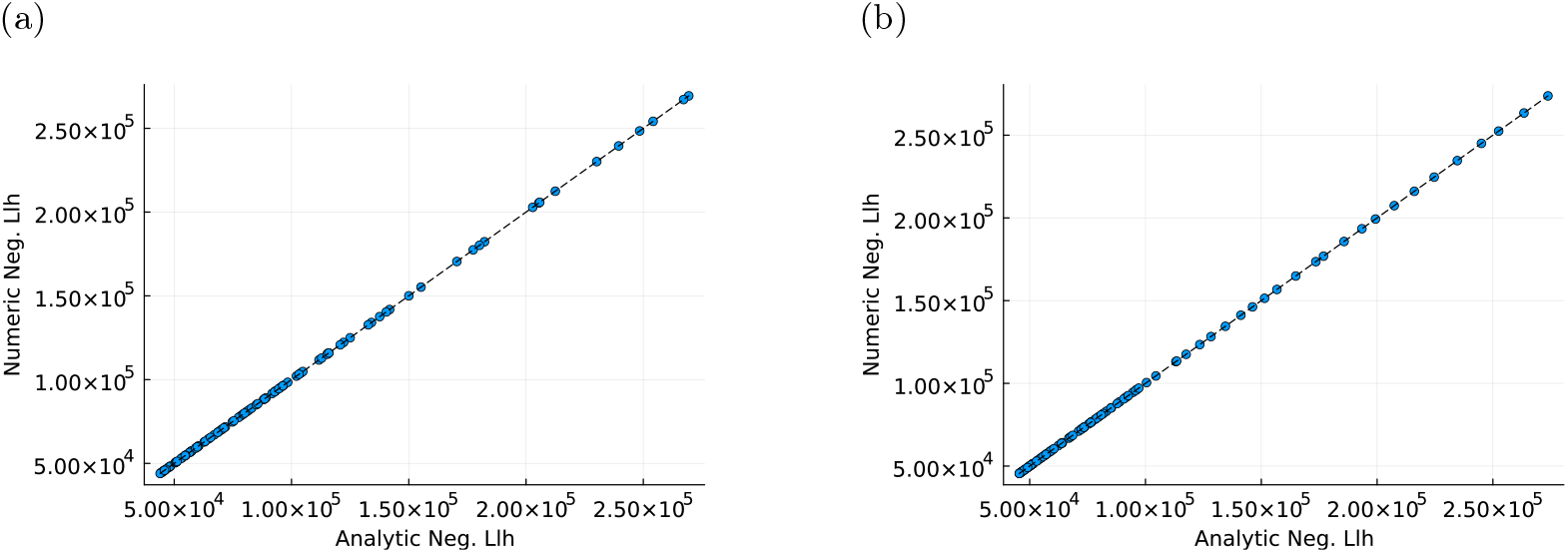
Comparison of likelihood formulations. We randomly sampled 100 parameter vectors and evaluated the analytical negative log-likelihood as well as the numerical negative log-likelihood on this vectors. 6a visualizes the resulting values for the exponential proportional model and 6b for the cell division model. The blackdashed lines correspond to the 45^*°*^ lines.

### 4.3 Efficiency Gain by Using Analytical Likelihoods

Next, we investigated the computation time for evaluating the analytical and numerical negative log-likelihood functions for the two exponential tumour growth based models (M1) and (M2). For this, we repeatedly evaluated each function on 100 randomly sampled parameter vectors and stored the mean computation time for each parameter vector.

For both models, evaluation of the analytical likelihood was computationally more efficient than for their numerically approximated counterpart (Figure 7). In the exponential proportional model, we even observe a more than fifty-fold speed-up. In the cell division model, the evaluation is around four times faster for the analytical likelihood. The precise mean computation times for the evaluation can be found in the Supplementary Information Table S3.

**Fig. 7:**
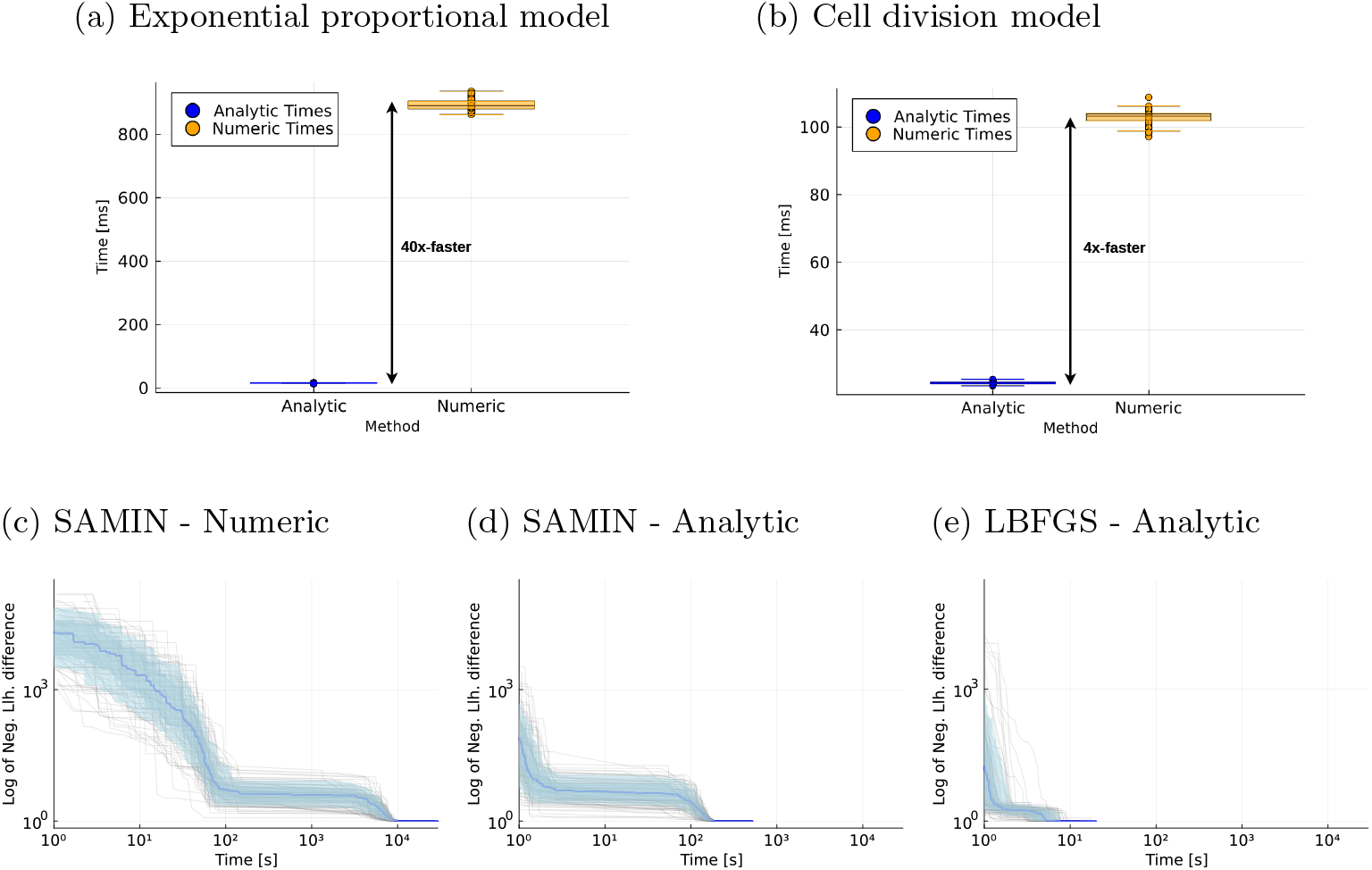
Evaluation of computational efficiency. Figures 7a and 7b show the mean computation times of evaluating the negative log-likelihood function on 100 randomly sampled parameter vectors for the two exponential growth based models (M1) and (M2). Figures 7c, 7c, and 7c depict the trace of the current best negative log-likelihood value over time for 100 single optimisation runs of the exponential proportional model using different likelihood formulations and optimization algorithms.

To assess the daily applicability of the combined model for analysing patient trajectories from clinical data, we are interested in the computation time of optimizing the negative log-likelihood of the models given the data. For each model we sampled 100 parameter vectors as start points for optimisation and in each optimisation run traced the current best negative log-likelihood value over time. We observed a four-time faster optimisation in the cell division model by using the analytical likelihood function instead of numerical approximations in the simulated annealing optimisation algorithm. Leveraging automatic differentiation, we computed gradients for the analytical likelihood functions, their second big advantage, and were able to use a gradient-based optimisation algorithm. This sped-up optimisation by another 100-times (Supplementary Figure S3). For the exponential proportional model computation time decreased fifty-fold by using analytical instead of numeric likelihoods. Using the gradient based optimiser for the local optimisation task, the efficiency gain was about another fortyfold, as depicted in Figure 7. In total, this resulted in a 2,000 times faster optimisation and shows the benefit of analytical likelihoods and their gradients whenever possible. The precise mean computation times for the optimisation runs can be found in the Supplementary Information Table S4.

Moreover, we observed that the likelihood formulation did not influence the capability of the optimiser to converge to the optimum and convergence of the runs shows great reproducibility of the optimisation (Supplementary Information Figure S7).

### 4.4 Analytical Likelihoods Facilitate Accurate Parameter Inference

The proposed modelling framework aims to identify rates that determine the disease dynamics for individual patients. Therefore, we need to assess the uncertainty of the parameter estimates. The uncertainties were computed using MCMC methods. We found that the credibility intervals cover the parameters used to generate the simulated data. For the exponential proportional model (M1) the true value lies even in the smallest credibility interval of 80% for three parameters, showing a high accuracy of the estimation procedure (Figure 8). The influence of the metastasis on the death process, given by *d*_metastasis_, has the highest variance, as the result of the dependence of this parameter on two stochastic processes. The highest relative deviation between true and estimated parameter is in the base rate of the metastatic spread given by *m*_basal_.

**Fig. 8:**
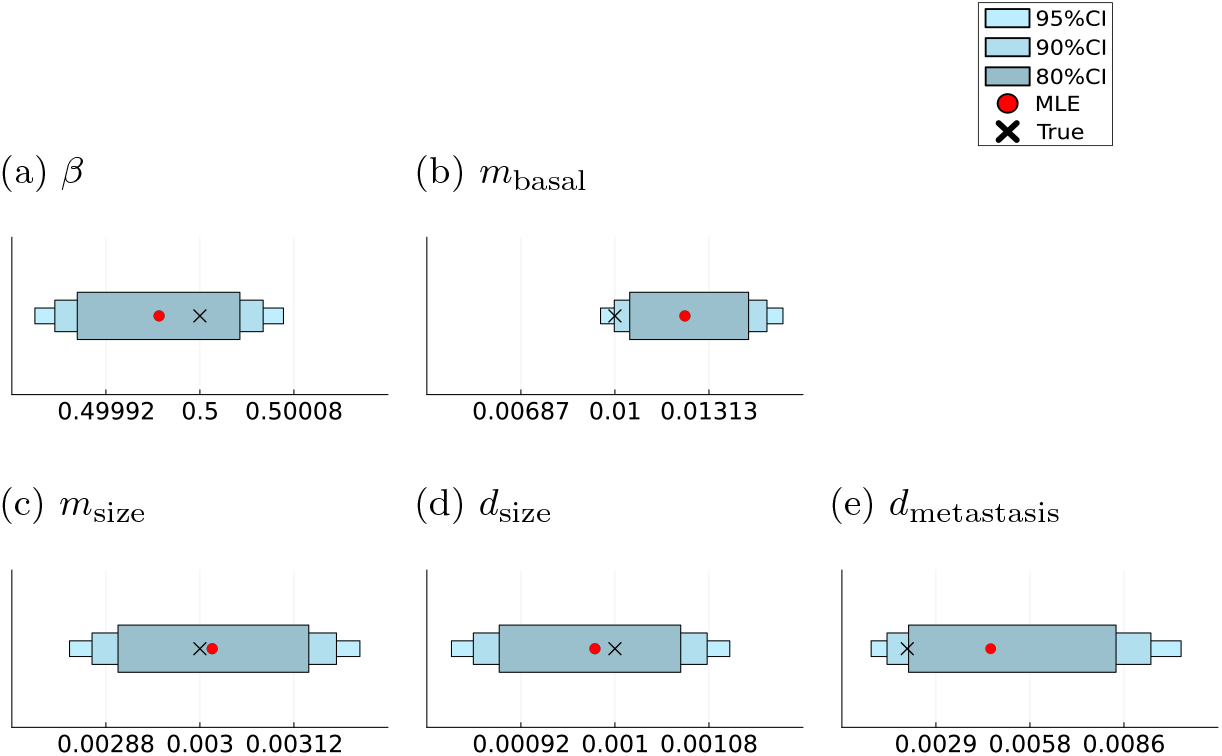
Parameter inference results for model (M1) Maximum likelihood estimates (MLE) and sampling-based credibility intervals for the model parameters of the exponential proportional model.

In the cell division model (M2), only one parameter *m*_division_ determines the trajectory of the metastasis process, reducing the uncertainty in the metastasis rate. Hence, for all four model parameters, the true values are covered not just by the 95%, but also by the 90% and 80% credibility intervals (Figure 9). This reveals the capabilities of the model to retrieve the underlying parameter values with high certainty. As before, the highest variance is in *d*_metastasis_ due to its stochastic nature. The high uncertainty in that parameter is also reflected in the estimated density of its marginal likelihood. In contrast to the other parameters, it admits a right-skewed distribution with a higher variance (Supplementary Information Figures S8, S9).

**Fig. 9:**
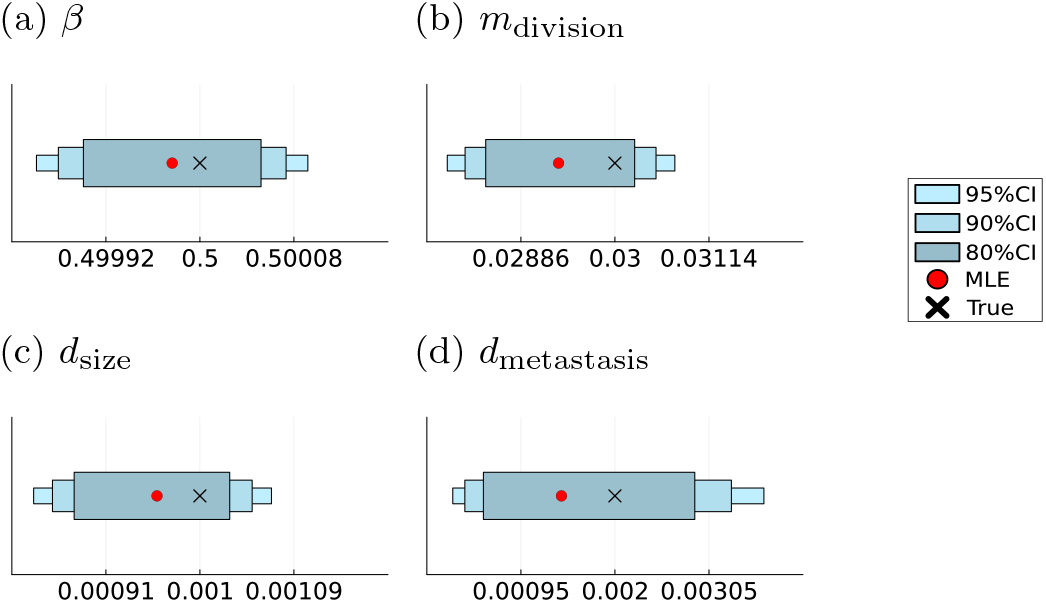
Parameter inference results for model (M2) Maximum likelihood estimates (MLE) and sampling-based credibility intervals for the model parameters of the cell division model.

In clinical application, the data at hand does not consist of equidistant and structured observations of the patients. To assess the performance of the proposed modelling and inference framework, we ran the optimisation for the exponential proportional model on a dataset including completely and partially missing timepoints. To be precise, we randomly removed about 30% of all measurement timepoints from the complete dense dataset with monthly observations. Additionally, we assigned about 20% of the tumour measurements and about 20% of the metastasis measurement to be missing. Using this sparse and dataset with partially missing timepoints, we still obtained accurate estimates ***θ***^*MLE*^ and credibility intervals covered the parameters used for data generation (Supplementary Information Figure S4). As expected the credibility intervals showed more uncertainty in the precise parameter values due to a less informative dataset.

### 4.5 Numerical Likelihood Approximations Enable Flexible Model Formulations

One further advantage of the proposed modelling framework is its flexibility. Yet, in the models considered so far, we only included models based on exponential growth, which allowed us to derive analytical likelihood formulations. However, the use of numerical integration to approximate the likelihood function allows for more flexible choices of the different processes. As examples for more complex tumour growth we consider the Gompertz growth model given in (3) and the Gyllenberg-Webb model based on equation (4). Both are combined with the volume based intensity rate function for the metastasis process as given in equation (5). For the Gompertz model we estimated the growth parameters ***θ***_Gompertz_[1, 2] = (*K, α*)^*T*^ and for the Gyllenberg-Webb model the proliferation and death rate ***θ***_GW_[1, 2] = (*b, µ*)^*T*^. All other parameters for the Gyllenberg-webb model were fixed to the example given in (Alzahrani et al 2014). We generated artificial data from the resulting models (M3) and (M4) using the model parameters provided in Table S7.

Subsequently, we performed 100 starts of minimizing the negative log-likelihood initialized at randomly sampled startpoints. For both models all starts converged to the same minimum (Supplementary Figure S11), yielding maximum likelihood estimators 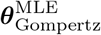 and 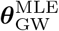 for the model parameters. Furthermore, the credibility intervals obtained by MCMC methods cover the parameter values used to generate the data as shown in Figure 10) and Figure 11. In the Gompertz model all parameters even lie in the the 90% credibility interval showcasing high certainty in the estimates. And again the highest variance is visible in the doubly stochastic parameter *d*_metastasis_. In the Gyllenberg-Webb model the base metastasis rate shows the highest uncertainty. However, all parameters are contained in the 95% credibility intervals.

**Fig. 10:**
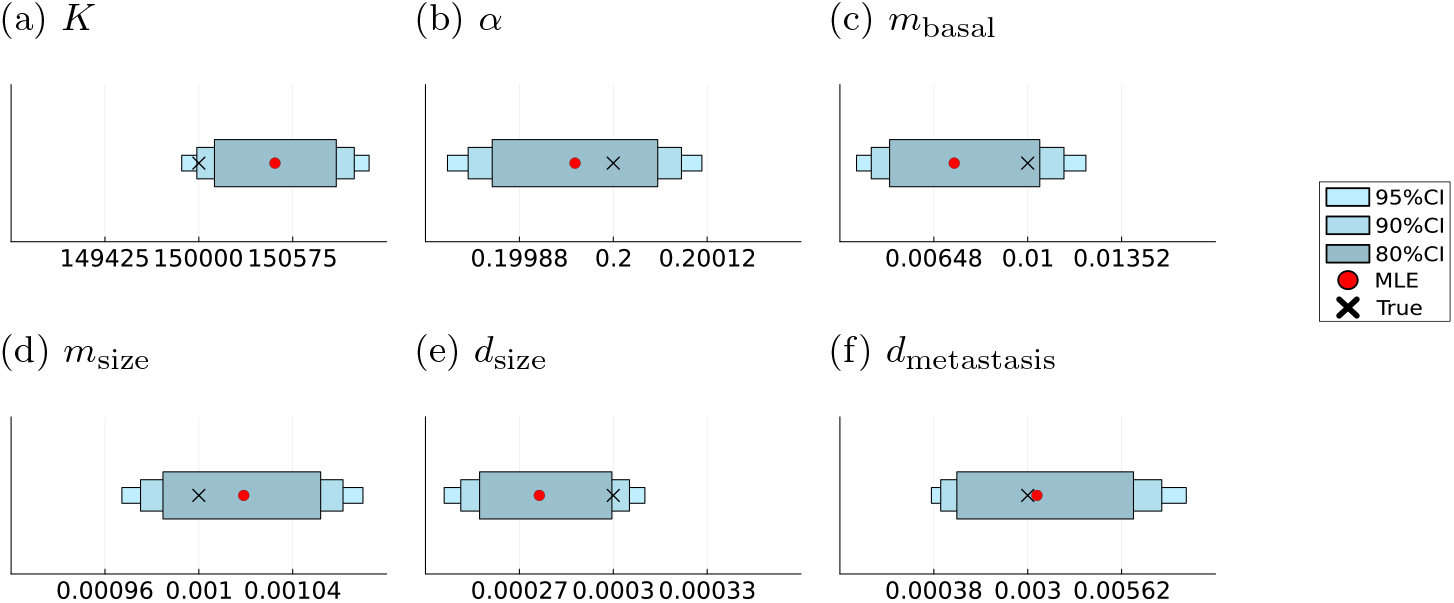
Parameter inference results for model (M3) Maximum likelihood estimates (MLE) and sampling-based credibility intervals for the model parameters of the Gompertz model.

**Fig. 11:**
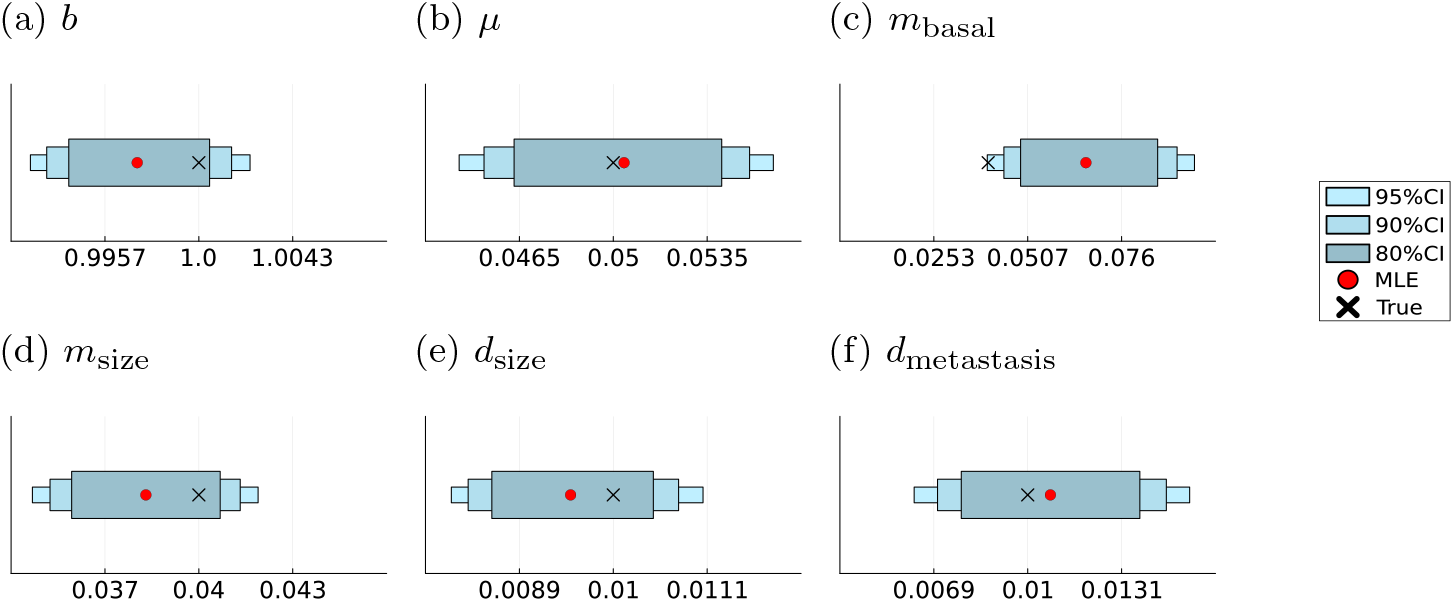
Parameter inference for model (M4) Maximum likelihood estimates (MLE) and sampling-based credibility intervals for the model parameters of the Gyllenberg-Webb model.

### 4.6 The Analysis Framework Allows to Distinguish Between Different Metastasis Processes

In biomedical studies the structure of the underlying processes is often partially unknown. In this cases, competing hypotheses can be compared using model selection. This can be achieved by comparing the Akaike information criterion (AIC) (Akaike 1974) of competing model formulations that try to explain the data. The AIC estimates the quality of each model by balancing between the goodness of fit and the simplicity of the model

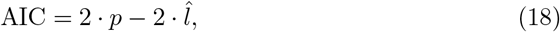

where *p* denotes the number of model parameter and 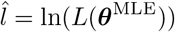 the maximum value of the log-likelihood function.

Here, we assess the ability of the proposed framework to determine the correct model structure, by creating artificial data for the two exponential tumour growth based models (M1) and (M2) (Figure 12) using parameters provided in Table S2. The metastasis processes differ in their dynamics and show a exponential like increase in the number of metastases for the exponential proportional model with volume based metastasis intensity function and a linear increase for the model with the cell division based metastasis intensity function (Figure 12).

**Fig. 12:**
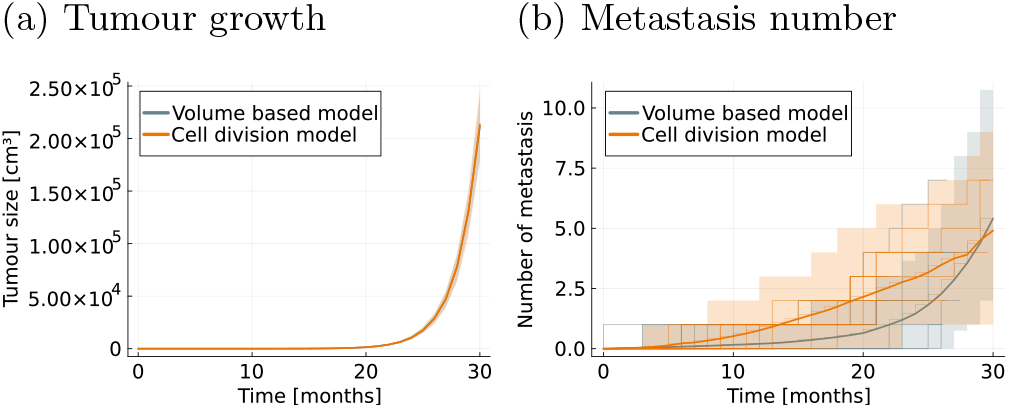
Datasets for model selection. Visual comparison of the two datasets used for the model selection task.

Each of the resulting datasets was subsequently fitted using both models the AIC values were computed. We observed that the AIC is minimal if the model coincides with the process underlying the data and we could then identify the correct model formulation. Furthermore, the difference in AIC value was significant for the wrong model formulation yielding high confidence in the selected model (Table 2). Similarly, the concordance between model and data is displayed when plotting the fit of observations against model simulations model obtained with the estimated parameter (Supplementary Information Figure S10).

**Table 2:**
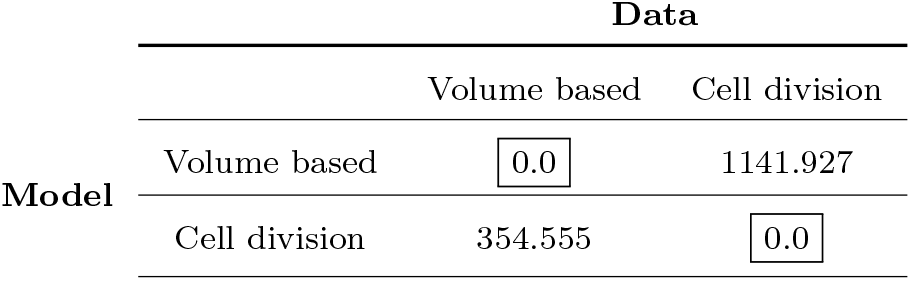
Model selection results. Difference of AIC values for the model given the dataset and the best AIC value obtained for that dataset.

### 4.7 The Model Supports Identification of Treatment and Covariate Effects

In addition to modelling individual patient trajectories, another goal in clinical research is the identification of treatment effects and covariates responsible for changes in the dynamics of the disease. Here, we extended the exponential proportional model by introducing a therapy effect that reduces the growth of the tumour as explained in Section 2.1.5. Additionally, we introduce the obesity status of the patient as a covariate that effects the therapy, where obesity is defined as a Body Mass Index greater or equal 30 (World Health Organization 2024). Obesity has been shown to be closely related to the development, characteristics and treatment efficacy of several cancer entities (Ross et al 2019; Wolin et al 2010). The realisation of treatment and obesity for a patient *l* are stored in the individual covariate vector

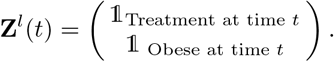

where we assume that the obesity status does not change over time and once a treatment is in place, it will be given until the end of the observations for that patient.

This results in the following functional form of the tumour growth rate

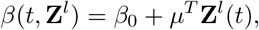

where *µ*^*T*^ = (−*ρ, δ*) is the vector of effects of the covariates on the basal growth rate *β*_0_ and encodes the presence of a treatment and the obesity status of the patient *l* at time *t*. We generated artificial data using parameters given in Table 3 in which the treatment start time and the obesity status were randomly drawn. Both covariates are assumed to be observed directly. We were able to reuse the computation of the analytical likelihood for the standard exponential proportional model to retrieve analytical likelihood for this model as well. Subsequently, we fitted the model running 100 starts of the LBFGS optimisation algorithm to obtain the MLE for the resulting model parameter vector ***θ*** = (*β*_0_, *ρ, δ, m*_basal_, *m*_size_, *d*_size_, *d*_metastasis_)^*T*^. All runs converged to the same optimal point (Supplementary Information Figure S13). The resulting estimates accurately recover the treatment effects and credibility intervals obtained by MCMC methods cover the parameters underlying the data (Figure 13). Opposed to the exponential proportional model the tumour growth parameters are not covered by the 80% credibility interval anymore, but by the 95% interval. However, the model still shows high certainty in this parameters shown by the small variance of these parameters and hence small credibility intervals.

**Table 3:** Treatment effect model parameterization. Model parameters with their corresponding ranges on logscale for the treatment effect model.

| Parameter | True value | Parameter range |
| --- | --- | --- |
| $\beta$ | -0693 | $[-0.75, -0.65]$ |
| $\rho$ | -0.799 | $[-0.85, -0.75]$ |
| $\delta$ | -1.609 | $[-2.3, -1.2]$ |
| $m_{\text{basal}}$ | -4.605 | $[-7, -2]$ |
| $m_{\text{size}}$ | -6.908 | $[-7, -2]$ |
| $d_{\text{size}}$ | -8.1117 | $[-9, -4]$ |
| $d_{\text{metastasis}}$ | -5.809 | $[-9, -4]$ |

**Fig. 13:**
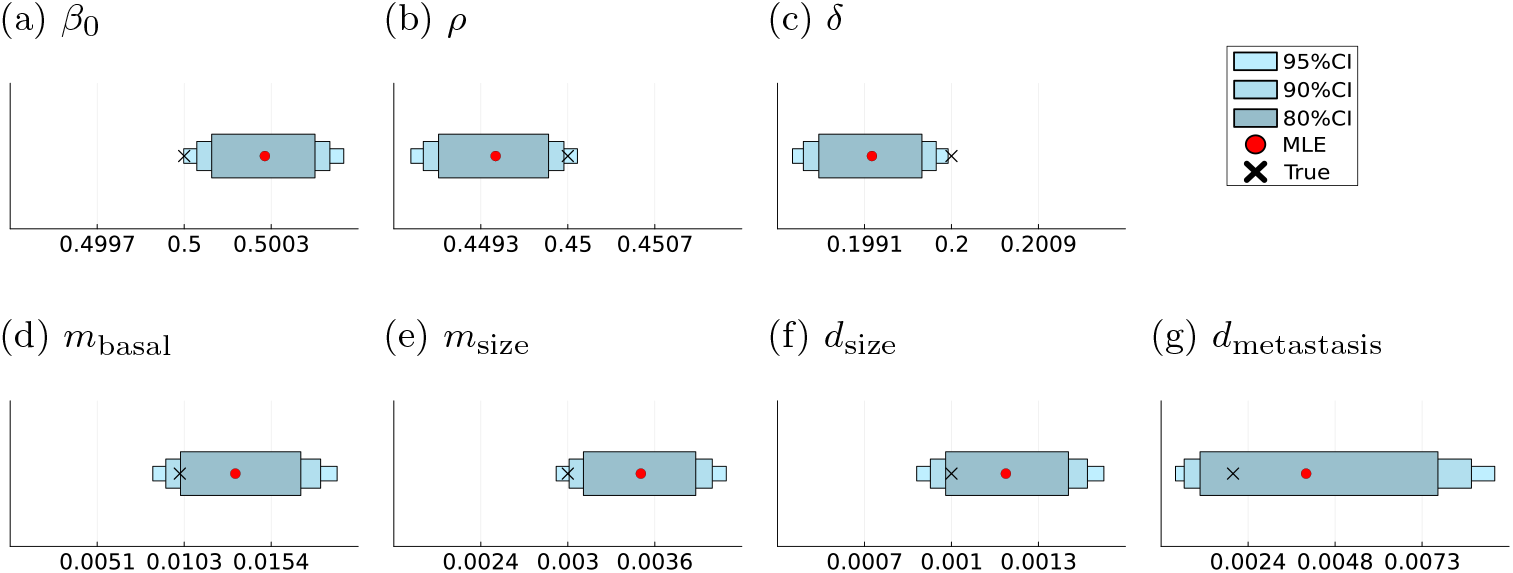
Parameter inference for model (M5) Maximum likelihood estimates (MLE) and sampling-based credibility intervals for the model parameters of the treatment effect model.

## 5 Discussion

The proposed class of combined stochastic models provide a useful alternative to multi-state or continuous growth models for studying cancer progression. While there exist many models that model tumour growth and metastatic seeding in special cancer entities, e.g. for breast cancer (Gasparini and Humphreys 2022), none of them directly couples this with the survival of the patients, or is flexible enough to be applied to different cancer entities. The modelling framework proposed in this paper explicitly combines the health status of the patient and the disease progression. Thus, it provides insights into the tumour growth and spread processes over time. It is the first of its kind and a first step towards holistic models for the analysis of patient trajectories and the development of practicable models for cancer progression.

The models we use in this work are examples of concrete model parameterisations. Yet, they still inherit some simplifying assumptions limiting their applicability in practice. In addition to the assumptions made in Section 2, we only applied them to simulated data, which allowed some simplifying assumptions on the initial point *S*_0_ and *t*_0_. When applied to real data, every patient would exhibit a different value for *S*_0_. However, given the model formulation, we can then compute the time until detection *t*_0_ and shift the time scale for each patient individually.

Moreover, the use of artificial data yields a more structured environment. We have shown that time intervals between observations need not to be equidistant and have unregular data sets. Therefore, we adapted the formulation of the likelihood to time points being unstructured and without complete observations, i.e. only some of the subprocesses are observed. Although, this does not fully resemble the complexity of data collected in clinical routine, it proves that the concept of the modelling framework can easily be applied to various realistic datasets.

In addition, although patients could differ in the realisation of their covariates in Section 2.1.5, leading to individual trajectories, we only considered the case where all model parameters are the same for all patients. This issue can be addressed by introducing random differences in the parameters across patients by for example the use of a mixed effect like description for the tumour growth rate.

Moreover, to take advantage of all the information contained in the patient trajectories, the processes involved in the model should include more covariates, such as biomarkers or general patient characteristics.Time-varying covariates can also account for resistance to cancer therapy over time. Additionally to introducing covariates on the cancer growth rates, one can also add covariates to the rates of the other subprocesses. Such adjustments are depending on the data and question at hand, but the proposed modelling framework can serve as a good exploratory tool. Therefore, future work on possible rate parametrizations and extensions of the likelihood formulations is encouraged.

Although, we consider here only simple models for the subprocesses, the proposed framework provides a high degree of flexibility. While our results demonstrate that already the considered process descriptions captured real-world data, model refinements which capture additional biological and clinical details should be explored, e.g., the study design and the training process to avoid model bias. The precise model formulation should depend on the prior knowledge about the cancer entity, the available data and the research question.

One simplifying assumption we made in this manuscript, is to only consider exponentially distributed survival times. As shown in Section 2.4 this does not limit the overall survival times to be exponentially distributed. However, further enhancements by considering more realistic survival time distributions, such as a Weibull or a Gamma distribution, can be of interest in ongoing research.

Another pathway of future extensions is to consider even more complex tumour growth and metastasisation models that can for example capture spatial dynamics and micrometastases. Using PDE based models for the tumour growth, such as the Greenspan model (Greenspan 1972; Bull and Byrne 2022), can be incorporated similarly to the physiologically structured Gyllenberg-Webb model (Gyllenberg and Webb 1990; Kuang et al 2018) by adapting the likelihood formulation. Altering the metastasisation process to a finer scale or even using a progression model for the tumour growth and metastasisation on a cellular level (Rupp et al 2024) can lead to more insights.

Our proposed modelling framework is mainly concerned with modelling the patient trajectories on a higher level using clinical routine data. We did combine processes acting on different scales by using a physiologically structured tumour growth model with an metastasis and survival process dependent only on the total tumour size. However, this does not consider the whole multi-scale nature of cancer yet. Models of this important aspect of cancer and for example account for micrometastases as in (L Rocha et al 2024) would need more fine grained data. Additionally, this would need further investigation on the formulation of the corresponding likelihood function and is out of the scope of this work. But we highly encourage further research in providing a combined multi-scale model.

While the flexibility of the modelling framework allows for building and simulation of such extended and more complex model parameterisation, another bottleneck lies in the efficiency of the parameter estimation procedure. If the formulated model exhibits an analytical likelihood formulation, maximum likelihood estimation can be done efficiently. Otherwise, approximating the likelihood or using likelihood-free and simulation based Bayesian inference schemes can lead to inefficient estimation tasks. Improvement of those for this class of models is out of the scope of this paper and a direction for following research.

In this paper, we leveraged automatic differentiation for the computation of gradients and gradient-based optimisation for the models based on an exponential tumour growth law. However, further investigation into the computation or numerical approximation of the likelihood can lead to differentiable objective functions for more models. This could then allow the use of gradient-based optimisation and a higher computational efficiency for those models as well. Depending on the model formulation and its observables, it is also possible to estimate parameters not jointly, but in a hierarchical manner. For example, in the cell division model with all three processes being observed, one could use the analytical and differentiable expressions for the tumour and metastasis rate parameters to estimate them with an efficient gradient-based method. The resulting estimates may then be fixed in the death process likelihoods, which are then optimized with the non-gradient based method. This can further improve the optimisation procedure. Hence, the inference procedure is a subject of ongoing research as well and could profit from further advances in computational frameworks and software packages.

One further direction for developing a more realistic description of cancer dynamics can be to evolve to a completely stochastic model by exchanging the deterministic growth laws for the tumour growth by their corresponding stochastic counterpart or a general SDE representation. More recent models suggest this exchange by stochastic differential equations for tumour growth (Ayuni Mazlan et al 2017; Mansour and Abobakr 2022; Katsaounis et al 2023). This then poses new challenges on computing or approximating the likelihood function and hence, the ability for efficient parameter inference. Therefore, further research on numerical approximations for such extended models or the use and efficiency of Bayesian estimation frameworks in this context can provide interesting new ways to extend this modeling framework further.

The combined stochastic model can address all six hallmarks of mathematical oncology (Bull and Byrne 2022). Moreover, the flexibility of the framework allows it to emphasize different aspects of the progression of cancer. As discussed before a tumour growth and metastasisation model could be used on a much more detailed scale, while keeping the survival process more general. Equivalently, a more detailed description of the survival process including tumour-immune interactions or immunotherapy can be used. Therefore, our framework is well suited to explore different model choices. These points may also be achieved by using agent-based models, they are in general computationally expensive and deriving a matching and complete set of decision rules can be challenging (Bull and Byrne 2023).

Even in the current form, our modelling framework provides a flexible starting point for exploring holistic models of cancer patients and enables future extensions that then accommodate more individualised stochastic processes and dependencies on various covariates such as therapy decisions. A modelling framework of this kind together with efficient inference is fundamentally important for studying cancer patient trajectories over time and providing practical value in clinical routine.

## Declarations

## Funding

This work was supported by the Deutsche Forschungsgemeinschaft (DFG, German Research Foundation) under Germany’s Excellence Strategy (EXC 2047—390685813, EXC 2151—390873048), by the German Federal Ministry of Education and Research (BMBF) under the CompLS program (GENImmune, grant no 031L0292F), by ERC grant INTEGRATE (grant no 101126146) and by the University of Bonn (via the Schlegel Professorship of J.H.).

## Conflicts of Interest

All authors declare that they have no conflicts of interest.

## Author Contributions Statement

**V.W**. Conceptualization, formulation of the mathematical model, computation of the likelihood functions, implementation of simulation, implementation of parameter estimation and uncertainty quantification, preparation of figures, writing the original draft, review and editing. **J.H**.: Conceptualization, formulation of mathematical model, review and editing of the manuscript, funding acquisition.

## A Supplementary Information

### A.1 Log-Normal Measurement Noise

In the simulation study shown in the main manuscript, we only considered Gaussian noise for the tumour size measurements. In general, the framework exhibits enough flexibility to deal with various noise formulations and the precise noise model that is used has to depend on the dataset at hand. To showcase this ability of our framework, we provide here the results for a experiment using the exponential proportional model together with log-normal distributed tumour size measurement noise, i.e.

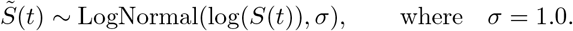

The optimization still showed good convergence, Supplementary Figure S14 and parameter estimates and corresponding credibility intervals covered the true values, highlighting the accuracy of the retrieved estimates, Supplementary Figure S15.

### A.2 Survival time distributions

Here we provide the parametrizations of the three survival time distribution as used in the manuscript.

#### Exponential distribution

The exponential distribution is the simplest and continuous analogue of the geometric distribution. It’s key property is the memoryless property which relates to having a constant hazard. We use the parametrization depending on a scale parameter *λ >* 0 and probability density function

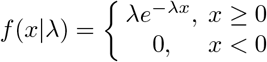

#### Weibull distribution

The Weibull distribution extends the exponential distribution by relaxing the assumption of constant hazard. Using a shape parameter *k >* 0 and a scale parameter *λ >* 0 the probability density function is given by

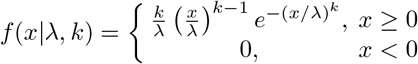

A value of *k <* 1 corresponds to a decreasing hazard over time and *k >* 1 to an increasing hazard over time, e.g. due to aging. In the case of *k* = 1 the Weibull distribution reduces to the exponential distribution with scale *λ*.

#### Generalized Gamma distribution

We use the parametrization with two shape parameters *d >* 0, *p >* 0 and a scale parameter *a >* 0 and support *x* ∈ (0, ∞). The probability density function is then given by

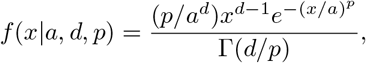

where Γ(·) denotes the gamma function.

This distribution generalizes the gamma distribution which has one shape parameter and also includes as special cases the exponential distribution, i.e. *d* = *p* = 1 and *λ* = *a*, and the Weibull distribution, i.e. *k* = *d* = *p* and *λ* = *a*.

## B Supplementary Figures

**Fig. S1:**
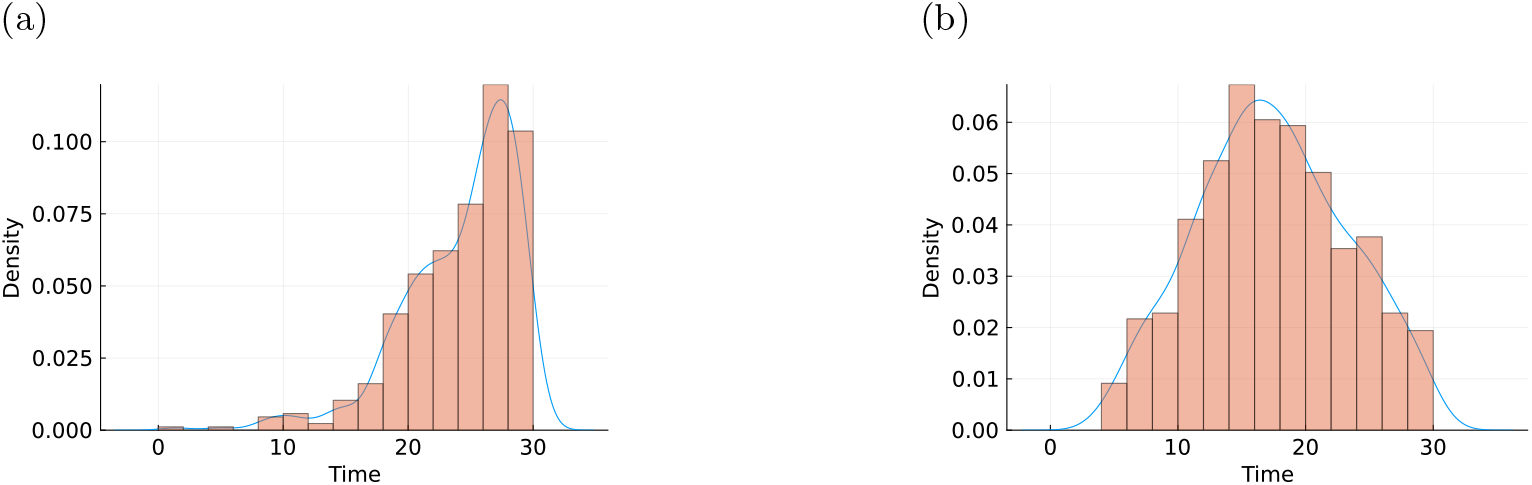
Overall survival time distributions. Histograms and kernel density estimates (blue lines) of the empirical distribution of overall survival times observed in the synthetic data sets for the exponential proportional model (M2) in S1a and the Gompertz model (M3) in S1b.

**Fig. S2:**
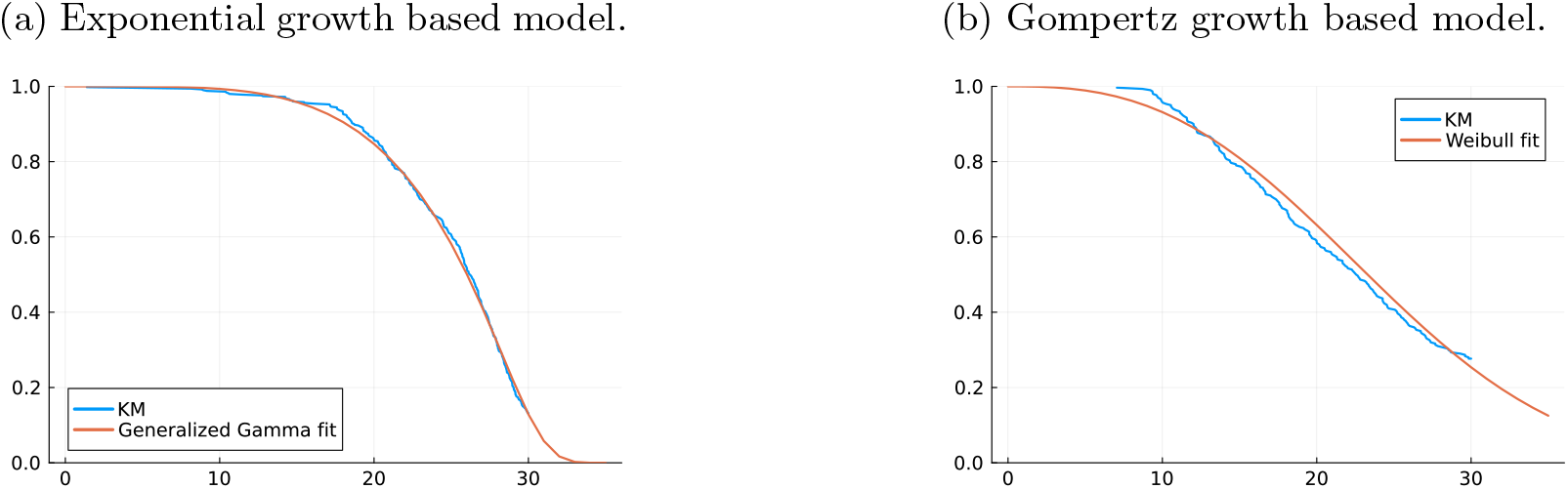
Assessment of survival curves. Kaplan-Meier estimator (KM) for the survival curve of the overall survival observed in the synthetic data (blue) plotted against the survival curve of a fitted distribution (orange). For S2a we used a generalized Gamma distribution with *a* = 4.5, *p* = 19.5, *d* = 30.9 and the exponential proportional model (M1), for S2b a Weibull distribution with *k* = 2.6, *λ* = 21.9 and the Gompertz model (M3).

**Fig. S3:**
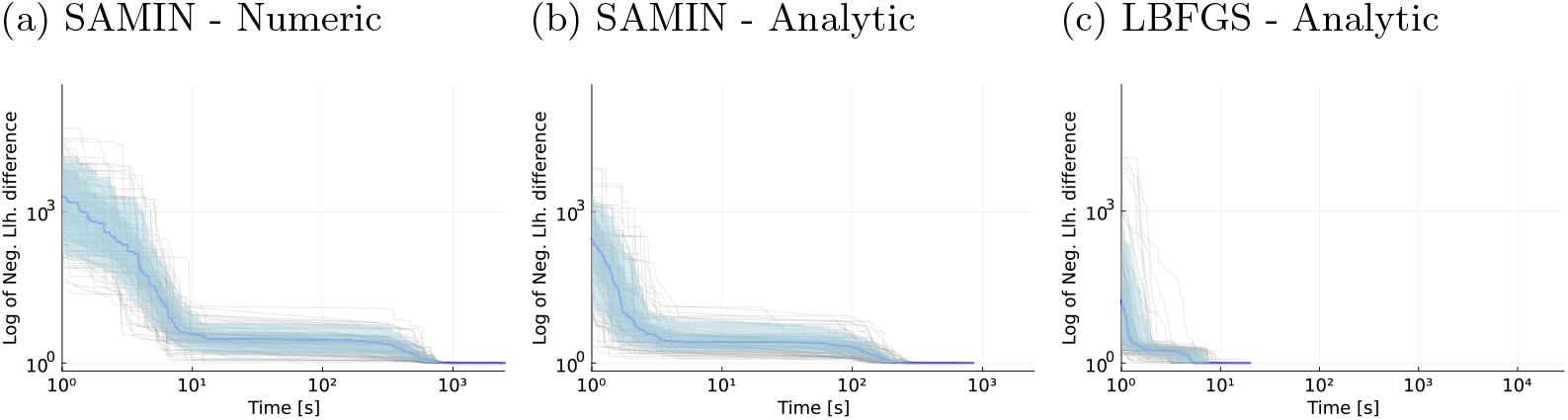
Optimization of the cell-division model. Optimiser traces of the current best negative log likelihood value over time for 100 single optimisation runs of the cell division model.

**Fig. S4:**
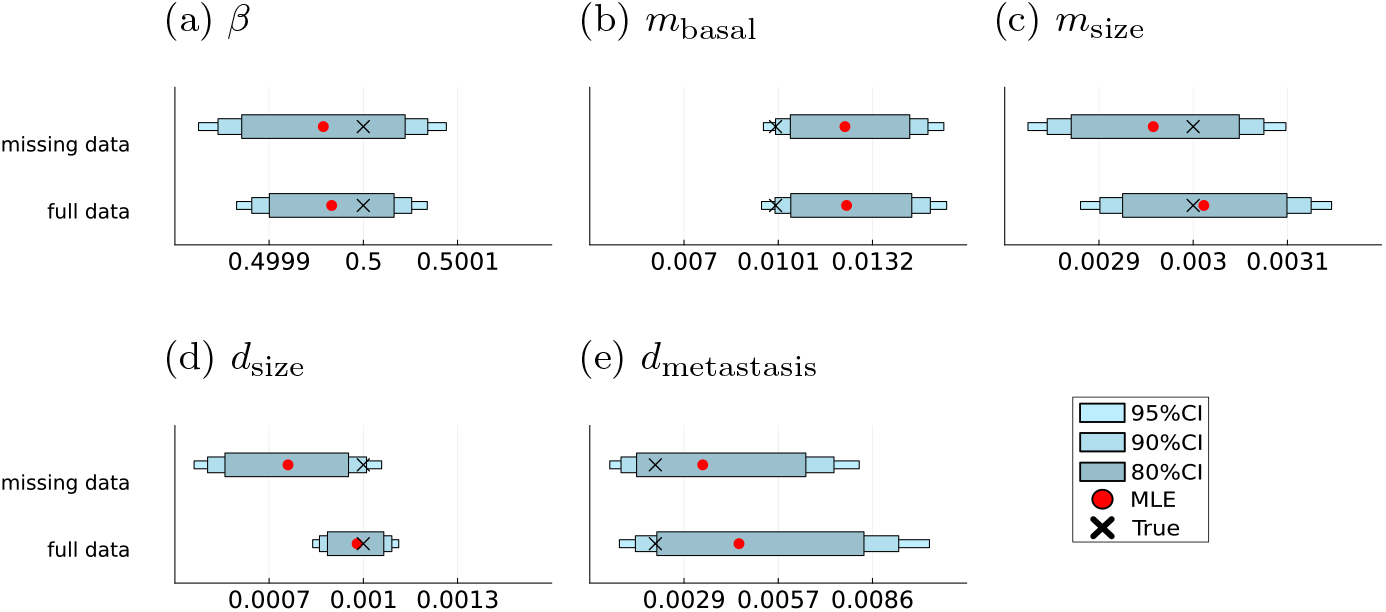
Parameter inference results for the exponential proportional model. Maximum likelihood estimates (MLE) and sampling-based credibility intervals for the model parameters of the exponential proportional model with the full data set and with an unregular data set, where we randomly subsampled 70% of the datapoints and added 20% of missingness into the tumour and the metastasis measurements each.

**Fig. S5:**
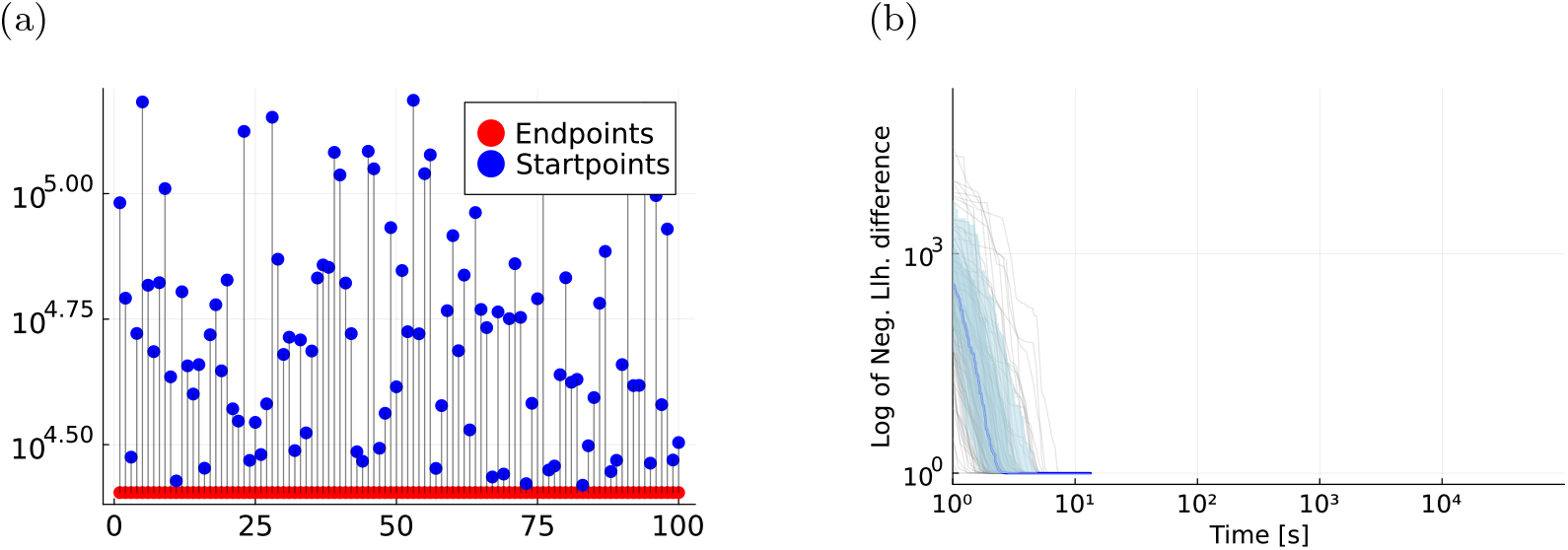
Optimization of the exponential proportional model with unregular data. Evaluation of 100 starts of LBFGS optimiser using analytical likelihoods and the exponential proportional model with an unregular data set, where we randomly subsampled 70% of the datapoints and added 20% of missingness into the tumour and the metastasis measurements each. S5a shows the double waterfall plot and S5b visualizes the optimizer traces.

**Fig. S6:**
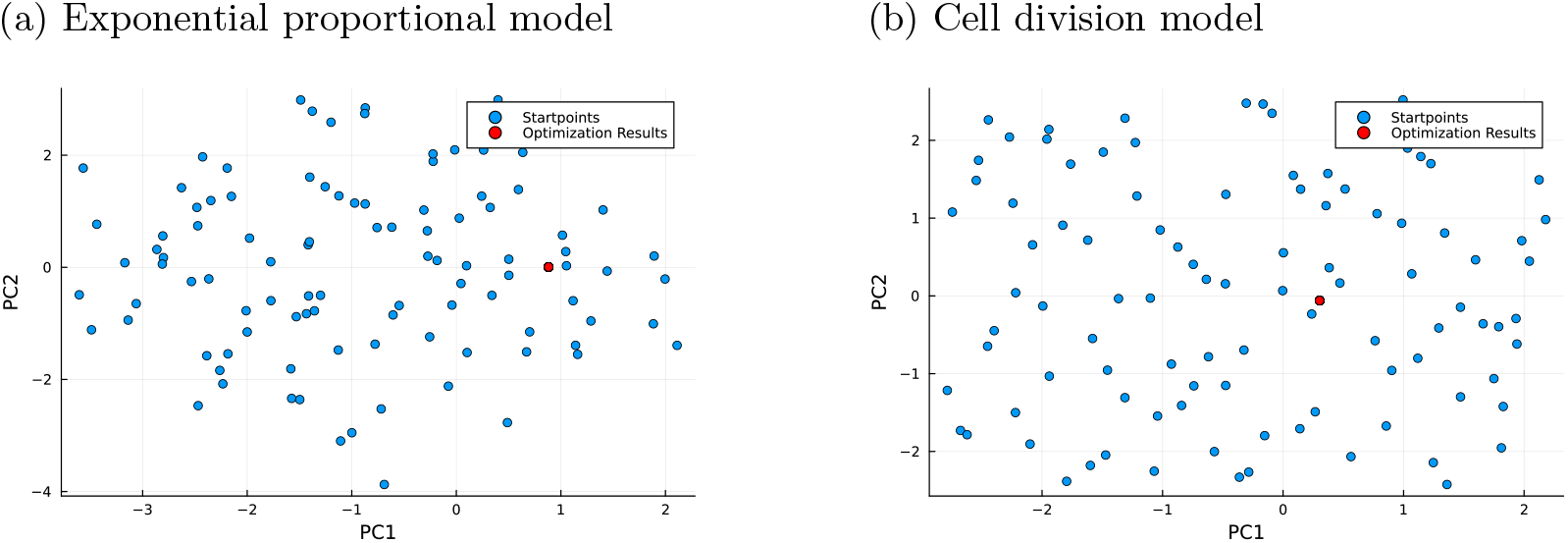
Illustration of optimization start- and endpoints. Visualization of the randomly sampled startpoints and the optimisation endpoint by a PCA on the parameter space.

**Fig. S7:**
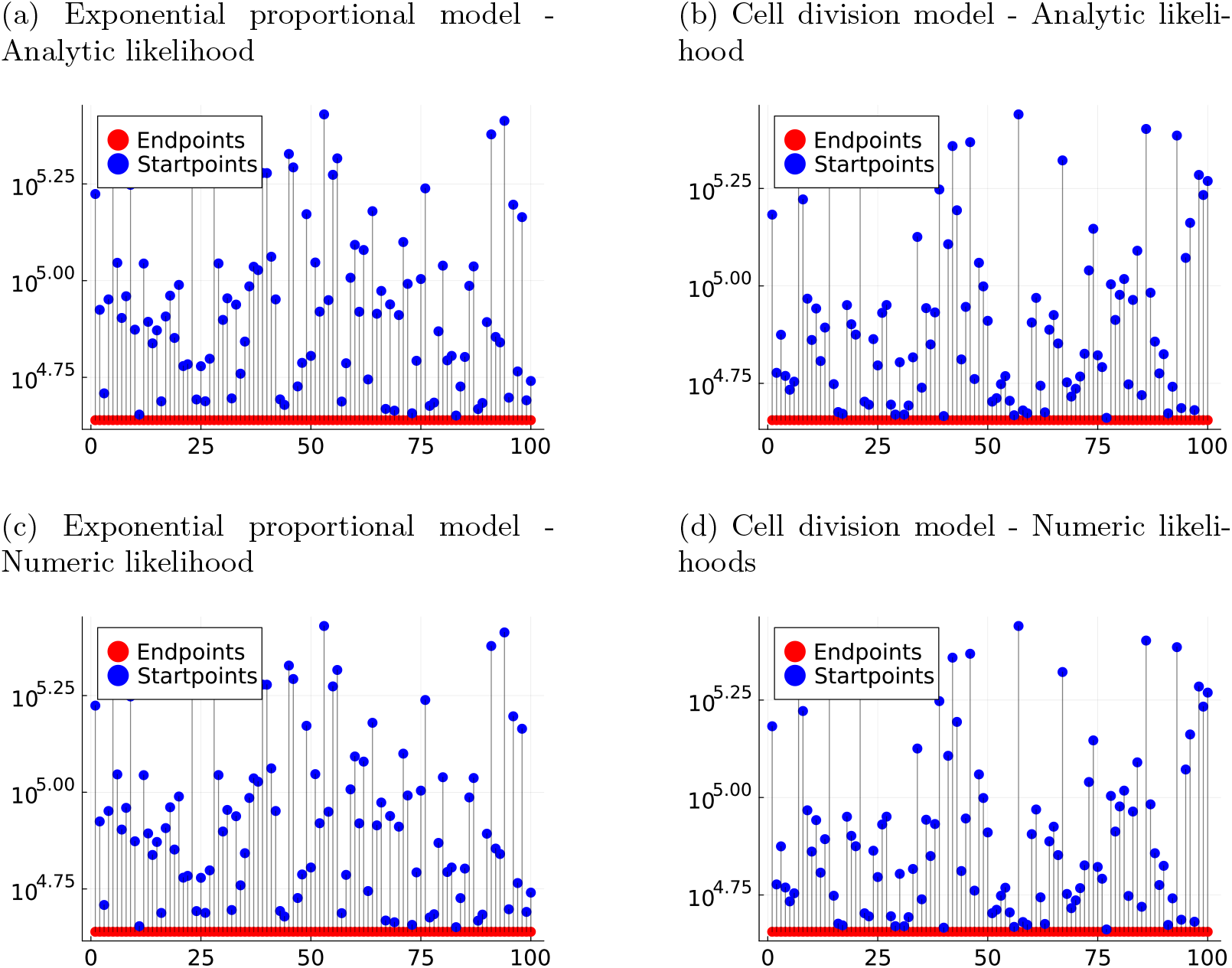
Convergence of SAMIN optimizations. Waterfall plots indicating the negative log-likelihood value of the startpoints and the corresponding endpoint of the optimisation. The plots where retrieved from the estimation results with analytical likelihoods and the SAMIN optimiser.

**Fig. S8:**
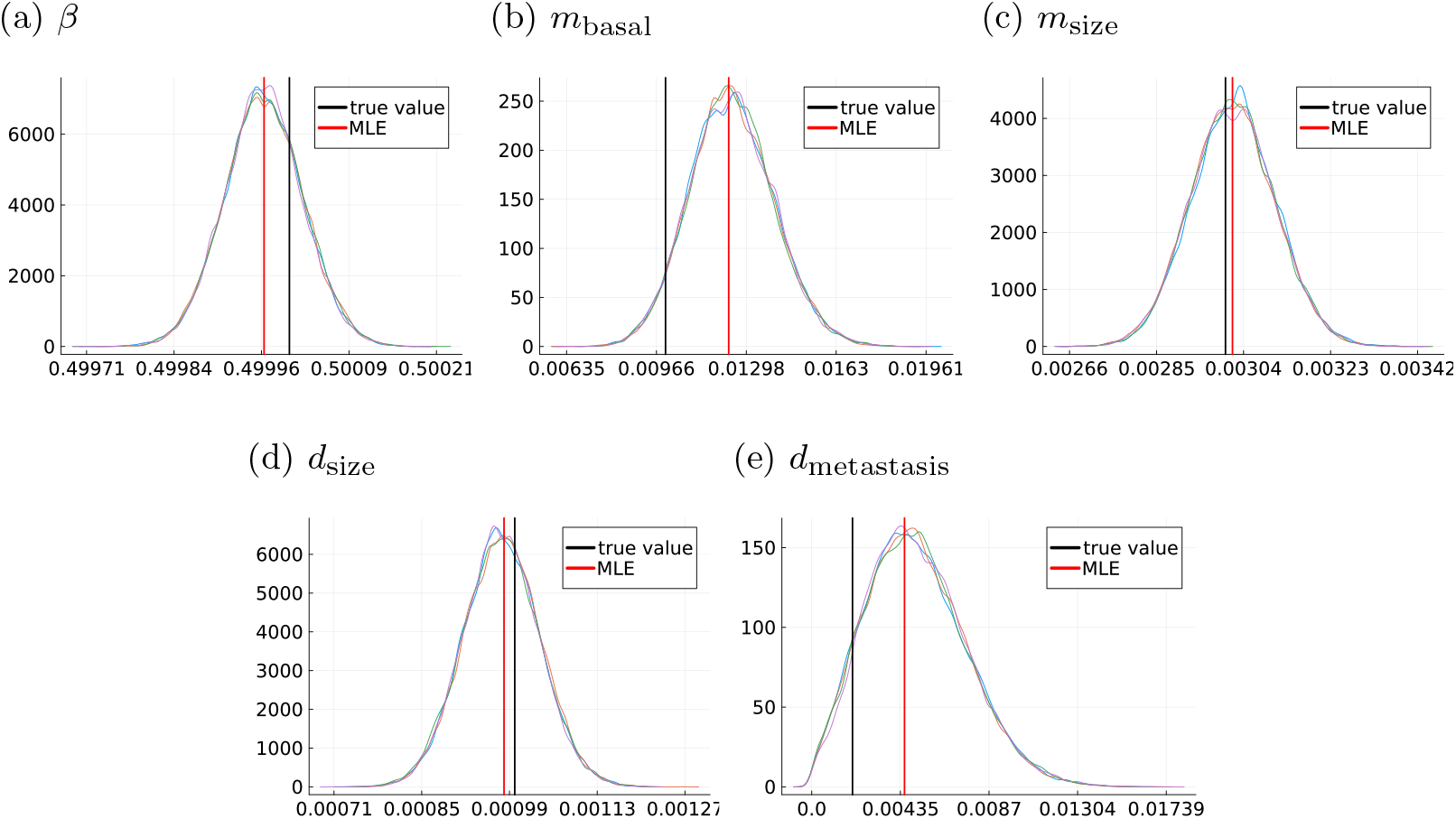
Sampling results of the exponential proportional model. MCMC sampling results visualized by density estimates over the samples for the model parameters of the exponential proportional model. The maximum likelihood estimator and the parameter value underlying the simulated data are indicated by vertical lines.

**Fig. S9:**
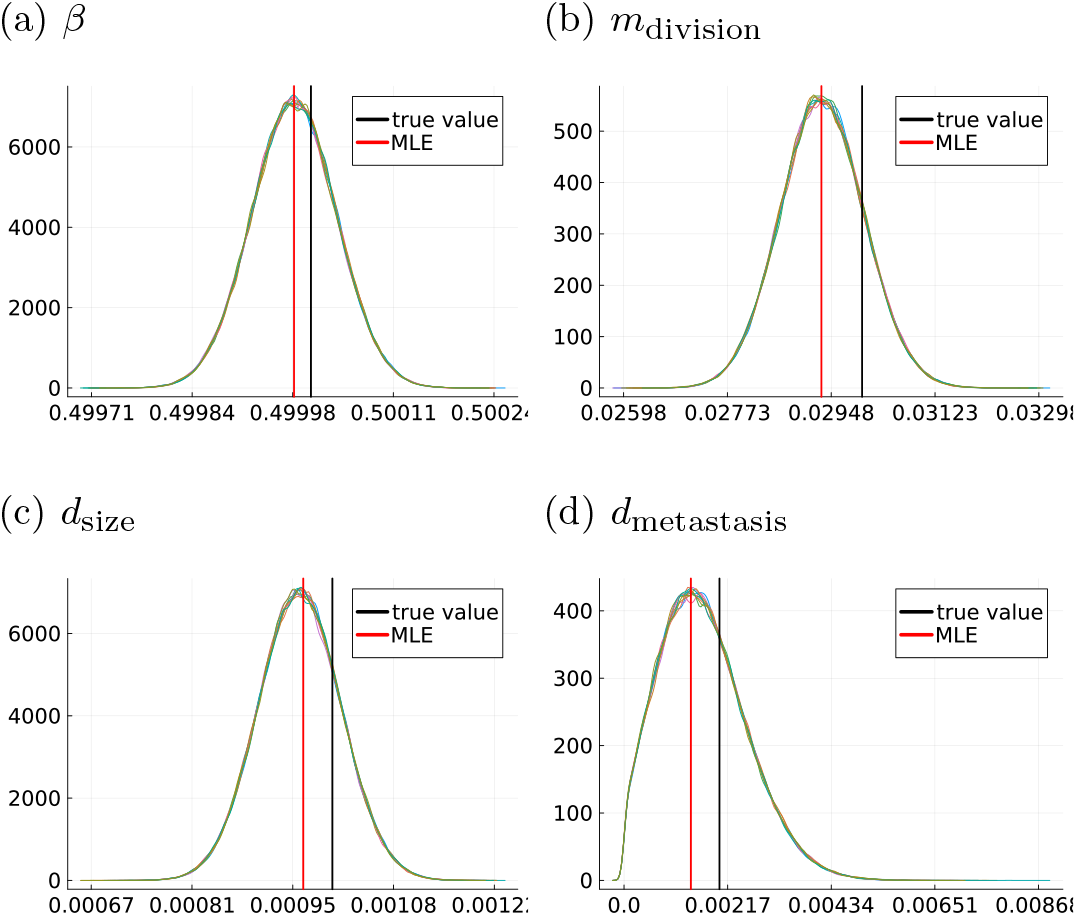
Sampling results of the cell-division model. MCMC sampling results visualized by density estimates for the model parameters of the cell-division model. The maximum likelihood estimator and the parameter value underlying the simulated data are indicated by vertical lines.

**Fig. S10:**
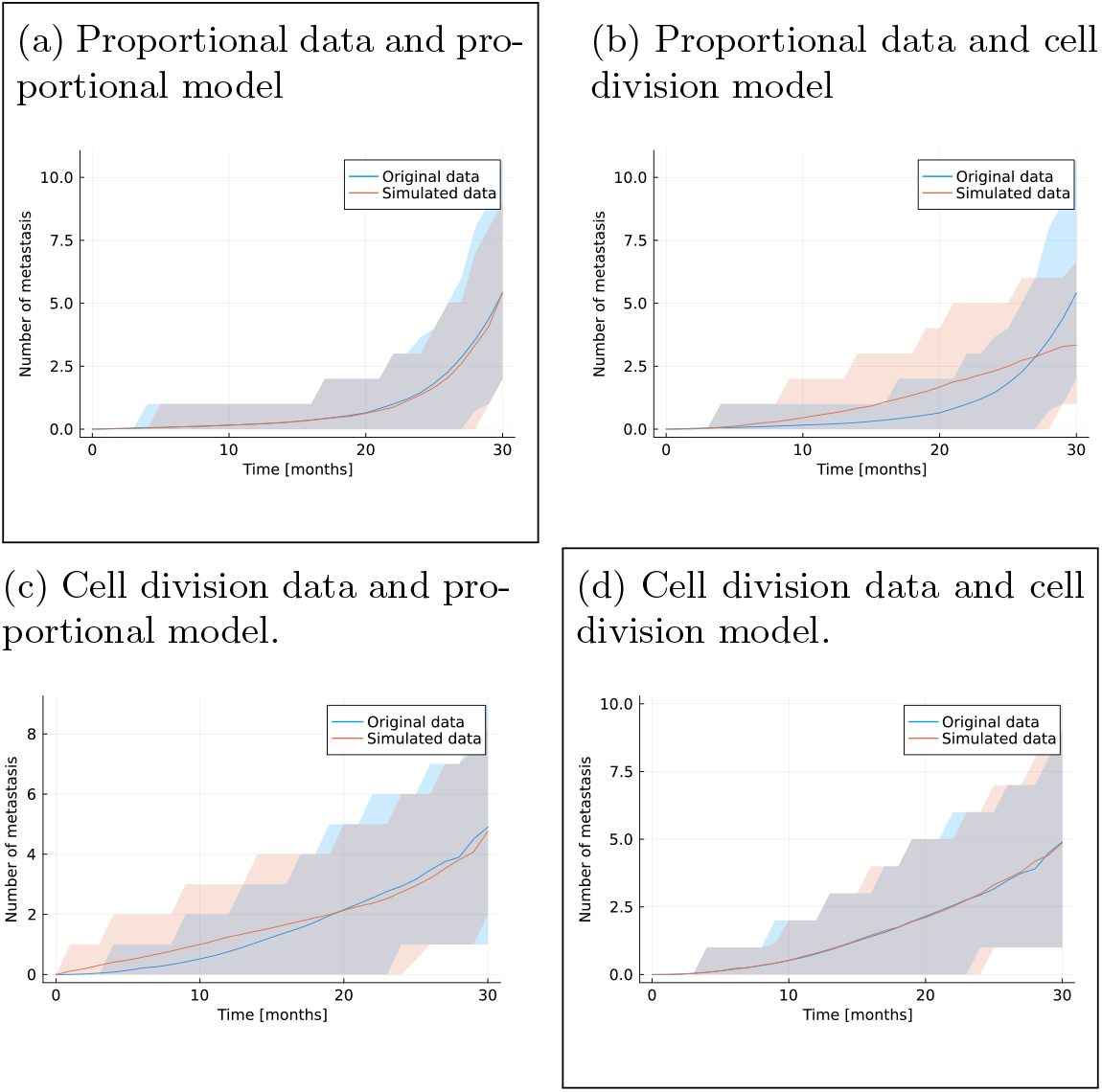
Model selection results. Model fits for the selection of the metastasis processes. We plotted the mean trajectory and a 95% confidence interval from the dataset used for the optimisation (blue) and for the data simulated with the model and corresponding MLE of the model parameters (orange). The boxes indicate the models with the smallest AIC value.

**Fig. S11:**
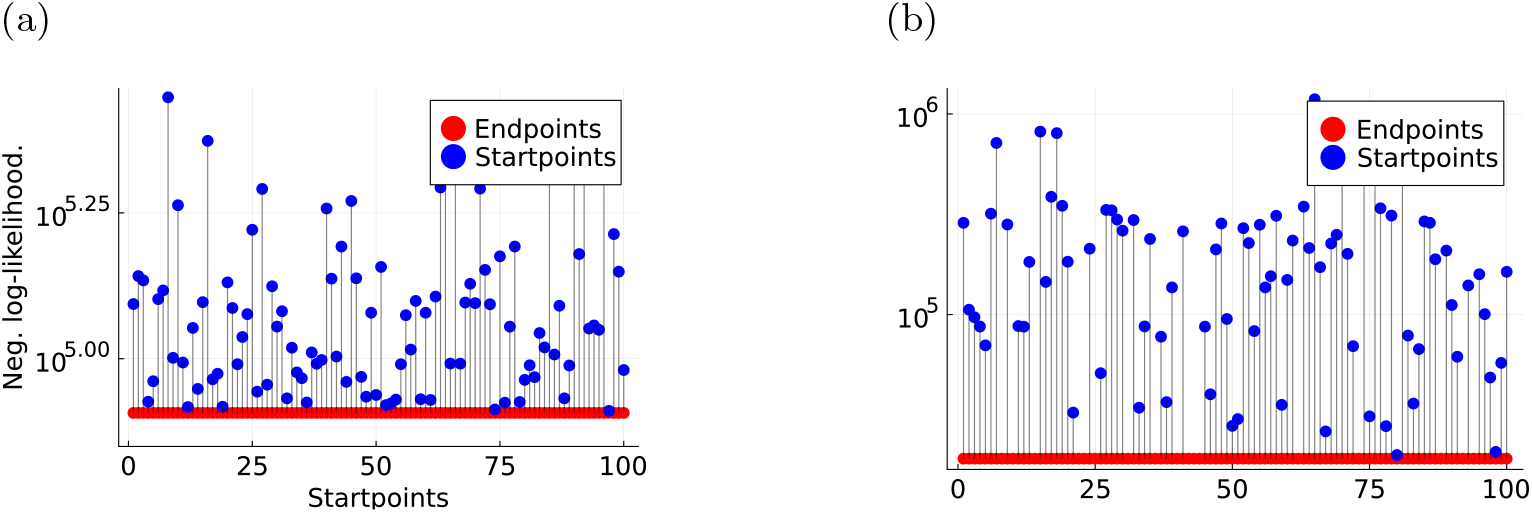
Double waterfall plot for the evaluation of the optimisation runs for S11a Gompertz model and S11b the Gyllenberg-Webb model. The plot shows the start- and endpoints of each optimisation run.

**Fig. S12:**
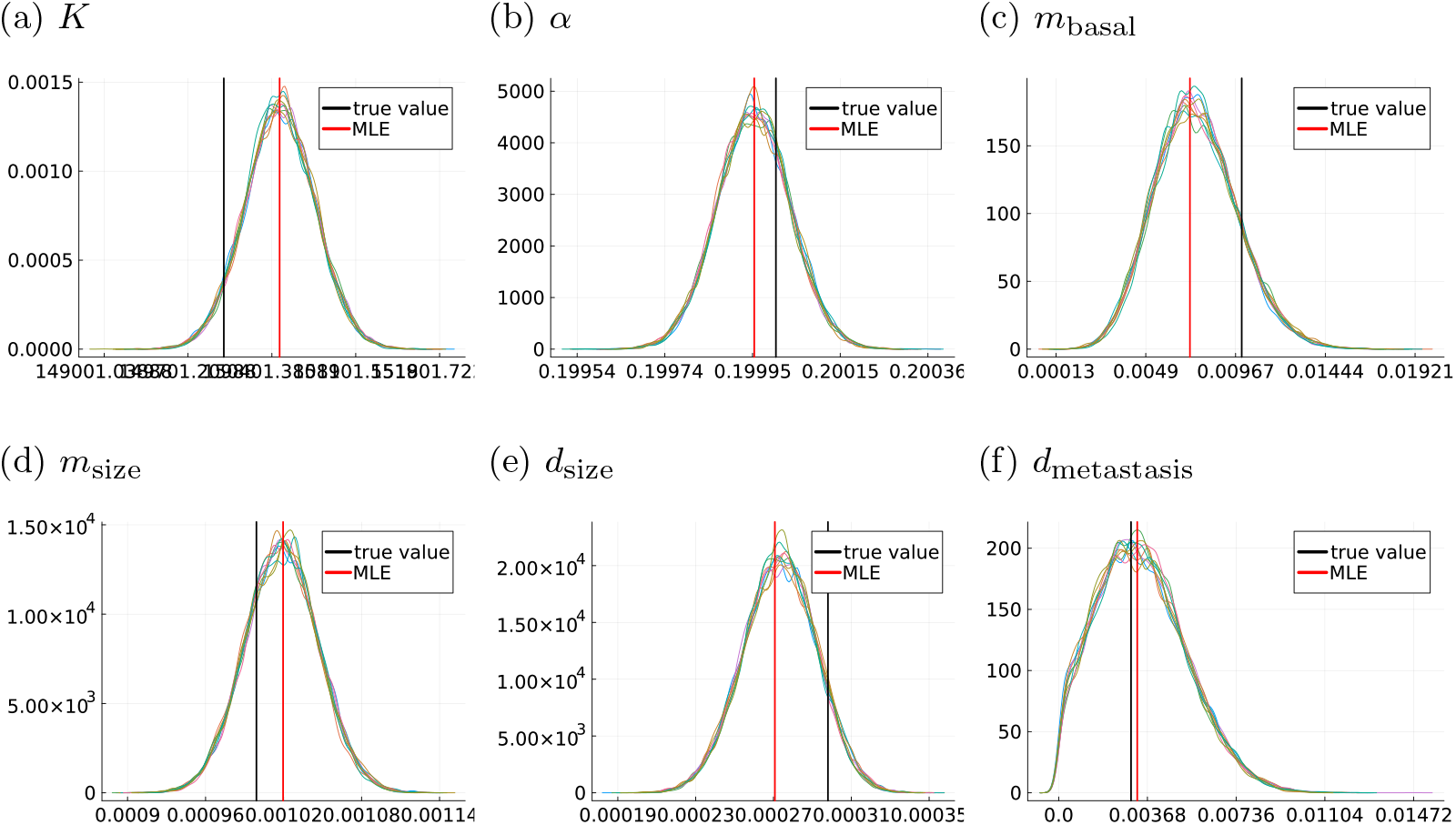
Sampling results of the Gompertz model. Estimation and sampling results visualized by density estimates for the model parameters of the Gompertz model. The maximum likelihood estimator and the parameter value underlying the simulated data are indicated by vertical lines.

**Fig. S13:**
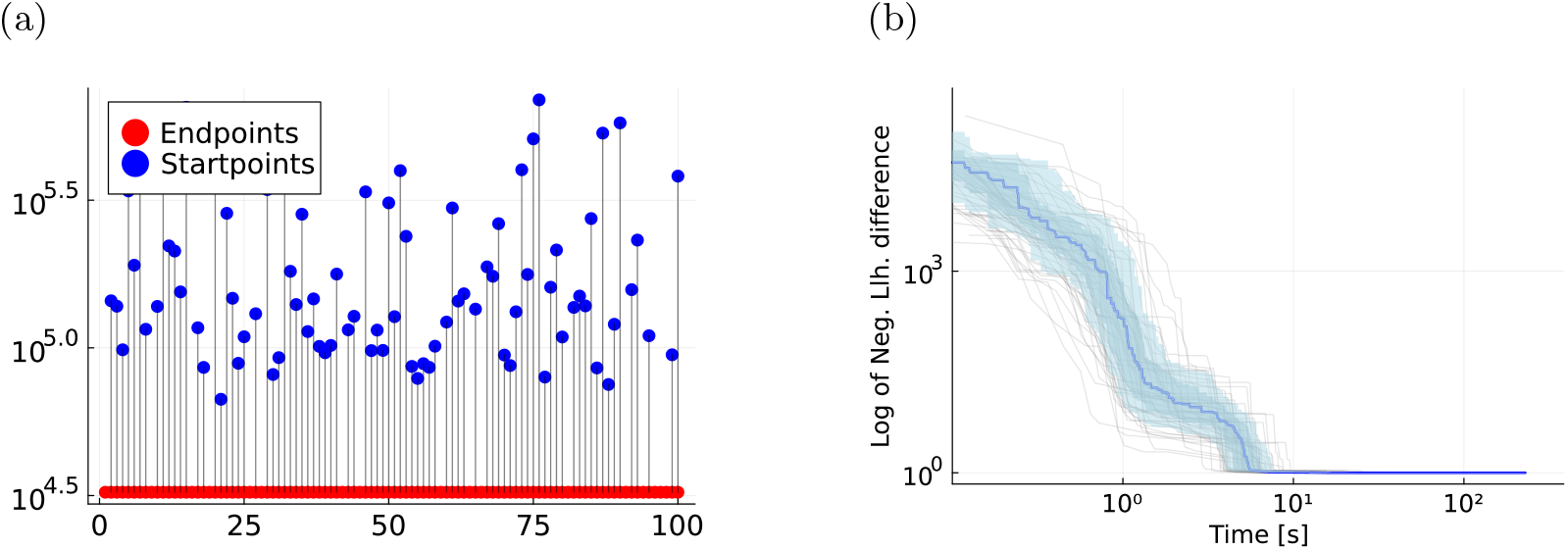
Optimization of the treatment effect model. Evaluation of 100 optimization runs of the treatment effect model. S13a shows the double waterfall plot and S13b visualizes the optimizer traces.

**Fig. S14:**
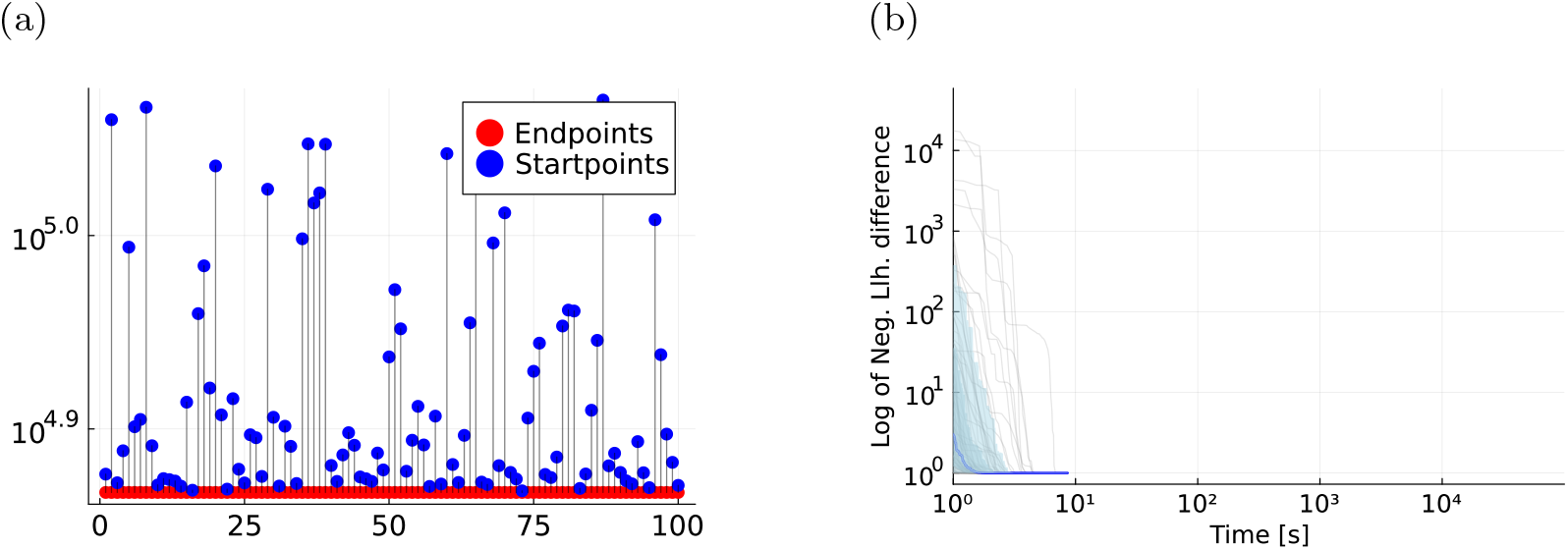
Optimization of the lognormal noise model. Evaluation of 100 starts of LBFGS optimiser using analytical likelihoods and the exponential proportional model with a lognormal noise model for the tumour size measurements. S14a shows the double waterfall plot and S14b visualizes the optimizer traces.

**Fig. S15:**
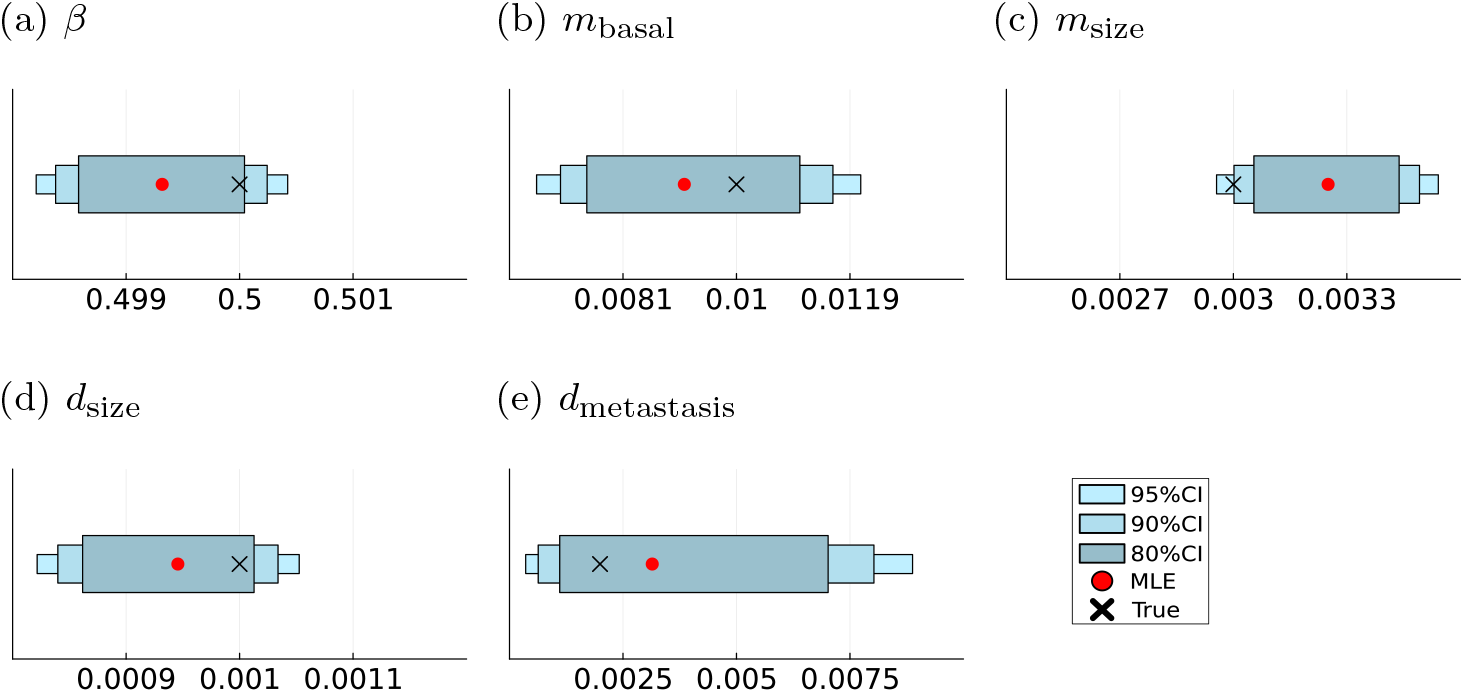
Parameter inference results for the lognormal noise model. Maximum likelihood estimates (MLE) and sampling-based credibility intervals for the model parameters of the exponential proportional model with log-normal measurement noise for the tumour size measurements.

## C Supplementary Tables

**Table S1:** Model parameters used in the model (M2) with exponential growth and cell division based metastasis spread in Section 2.4 to mimick the data characteristics from Engel et al (2003).

| Parameter | Value |
| --- | --- |
| $\beta$ | 0.01 |
| $m_{\text{division}}$ | 6.0 |
| $d_{\text{size}}$ | 0.011 |
| $d_{\text{metastasis}}$ | 0.31 |

**Table S2:** Model parameters, their true values and respective ranges for exponential growth based models on log-scale.

| Parameter | True value | Parameter range |
| --- | --- | --- |
| $\beta$ | -0.693 | $[-0.75, -0.65]$ |
| $m_{\text{basal}}$ | -4.605 | $[-7, -2]$ |
| $m_{\text{size}}$ | -5.809 | $[-7, -2]$ |
| $m_{\text{division}}$ | -3.506 | $[-7, -2]$ |
| $d_{\text{size}}$ | -6.908 | $[-9, -4]$ |
| $d_{\text{metastasis}}$ | -6.215 | $[-9, -4]$ |

**Table S3:**
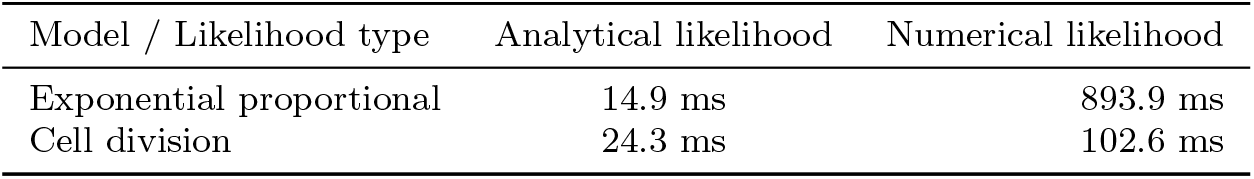
Mean evaluation times for the function evaluation of the negative log-likelihood function. The mean was taken over the evaluation times of 100 randomly sampled parameter vectors.

**Table S4:** Mean run-time of the optimisation. The mean was taken over 100 optimisation runs initialized at 100 randomly sampled startpoints.

| Model / Likelihood type and optimiser | Analytical - LBFGS | Analytical - SAMIN | Numerical - SAMIN |
| --- | --- | --- | --- |
| Exponential proportional | 13.3 s | 501.1 s | 26565.6 s |
| Cell division | 5.8 | 684.4 s | 2375.1 s |

**Table S5:** Exponential proportional model estimation results.

| Parameter | True value | Estimate (95%-CI) |
| --- | --- | --- |
| $\beta$ | 0.5 | 0.4999 (0.4998; 0.5) |
| $m_{\text{basal}}$ | 0.01 | 0.009147 (0.0067; 0.0125) |
| $m_{\text{size}}$ | 0.003 | 0.0029 (0.0027; 0.0031) |
| $d_{\text{size}}$ | 0.001 | 0.00097 (0.0008; 0.0011) |
| $d_{\text{metastasis}}$ | 0.002 | 0.0035 (0.0003; 0.0103) |

**Table S6:**
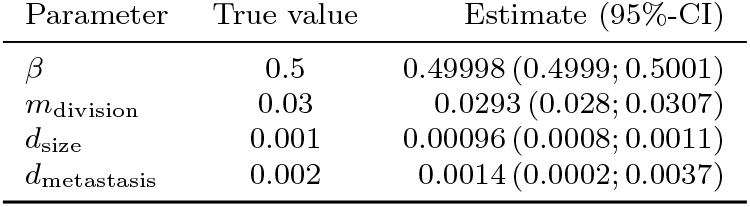
Cell division model estimation results.

**Table S7:**
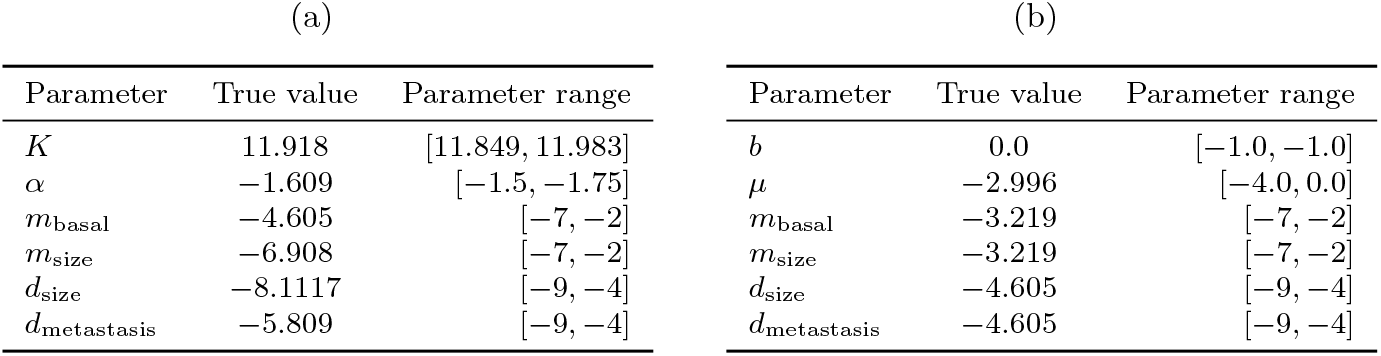
Model parameters with their corresponding ranges on log-scale for the Gompertz model S7a and the Gyllenberg-Webb model S7b.

(a)
| Parameter | True value | Parameter range |
| --- | --- | --- |
| $K$ | 11.918 | [11.849, 11.983] |
| $\alpha$ | -1.609 | [-1.5, -1.75] |
| $m_{\text{basal}}$ | -4.605 | [-7, -2] |
| $m_{\text{size}}$ | -6.908 | [-7, -2] |
| $d_{\text{size}}$ | -8.1117 | [-9, -4] |
| $d_{\text{metastasis}}$ | -5.809 | [-9, -4] |

| Parameter | True value | Parameter range |
| --- | --- | --- |
| $b$ | 0.0 | [-1.0, -1.0] |
| $\mu$ | -2.996 | [-4.0, 0.0] |
| $m_{\text{basal}}$ | -3.219 | [-7, -2] |
| $m_{\text{size}}$ | -3.219 | [-7, -2] |
| $d_{\text{size}}$ | -4.605 | [-9, -4] |
| $d_{\text{metastasis}}$ | -4.605 | [-9, -4] |

## D Proofs

In this supporting section we give the proofs for Theorem 1 and Corollary 2.

**Proof of Theorem 1**

*Proof*. For notational simplicity we do not explicitly write the dependence on other processes for the intensity rates and write *λ*_*N*_ (*t*), Λ_*N*_ ([*t*_*j−*1_, *t*_*j*_)), *λ*_*D*_(*t*), Λ_*D*_([*t*_*j−*1_, *t*_*j*_)). Additionally, we denote with Λ_*D*_([*t*_*j−*1_, *t*_*j*_), *n*) the accumulated death process intensity function for a time-interval, where the number of metastasis *n* is constant.

Let us first consider the simpler case of *m* = 0 new metastasis in the time interval [*t*_*j−*1_, *t*_*j*_) and *D*(*t*_*j*_) = *D*(*t*_*j−*1_) = 0, the patient survived during the time interval of interest. Then by the property of exponentially distributed waiting times in a Poisson process, the likelihood contribution is given by.

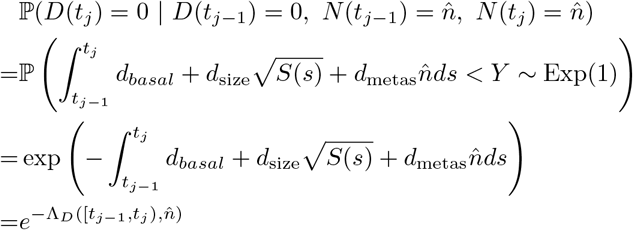

For shorter notation we name the solution of the last integral by

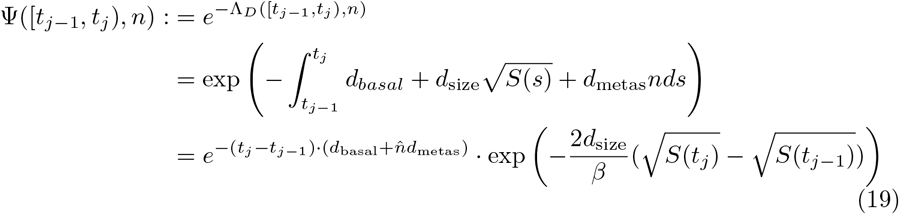

If we observe *m* = 1 new metastasis in the time interval of interest, we can simply split the interval at that timepoint *u*, where the metastasis occurred and get

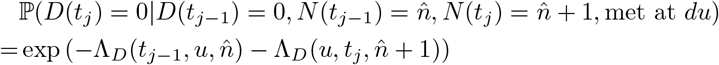

However, since we do not observe this timepoint, we need to integrate over the full time interval with respect to the probability that the metastasis occurred at that timepoint. This probability is given by

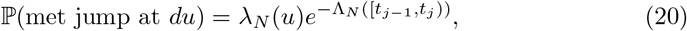

where *λ*_*N*_ (*u*) denotes the instantaneous rate of a jump at *u*

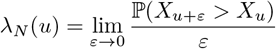

and 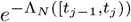 is the normalization constant based on the mean of jumps in the interval^4^

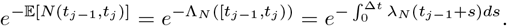

However, this omits the information that we already conditioned on having exactly one jump in the interval, so we need to divide by that probability.

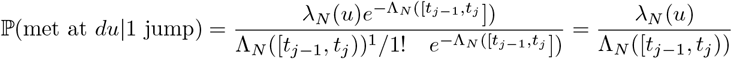

Together this yields:

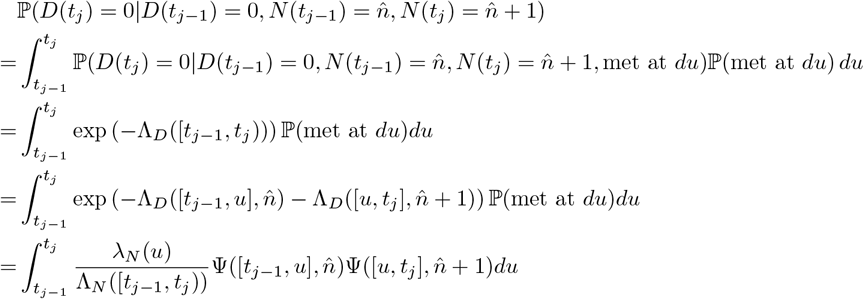

Analogously, we get for the case of *m* new metastasis, 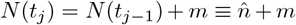.

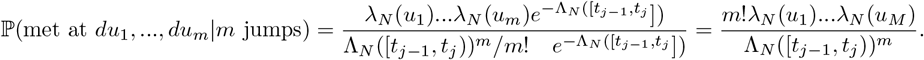

This yields the desired result

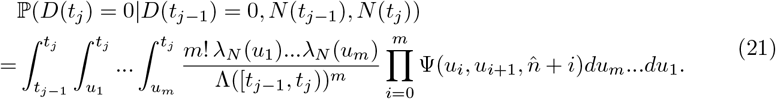

**Proof of Corollary 2**

*Proof*. For the case *D*(*t*_*j*_) = 1, *D*(*t*_*j−*1_) = 0, we now that death occurs exactly at time *t*_*j*_. So, we need to get the density for the time until an event given we did not see it before.

By the definition of a non-homogeneous Poisson process we know

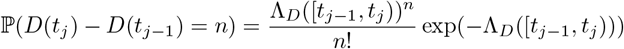

Therefore, we get for the distribution function for the time *T* until the next event after *t*_*j−*1_

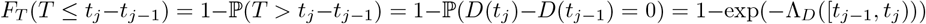

Differentiating this yields as a density for the time to next event after *t*_*j−*1_

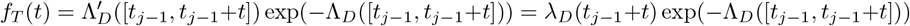

Intuitively this is the probability to survive until *t*_*j−*1_ + *t* multiplied with the instantaneous rate of dying at that time.

We then apply Lemma 1 for the form of the survival probability, given that *m* new metastasis occurred,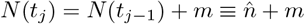. This then yields

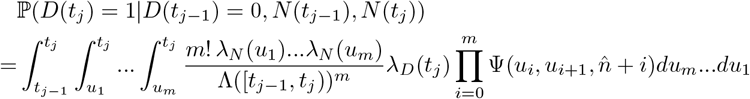

## E Analytical Likelihood Formulas

In this supplementary part, we provide the precise expressions of the analytically computed likelihood contributions (14) used in the simulation studies in Section 4. We incorporate the assumption of *t*_0_ = 0 and make the dependence on parameters explicit in the arguments of the functions.

### Cell Division Model

Given the following functions for tumour growth and intensity rates of the Poisson point processes

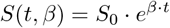

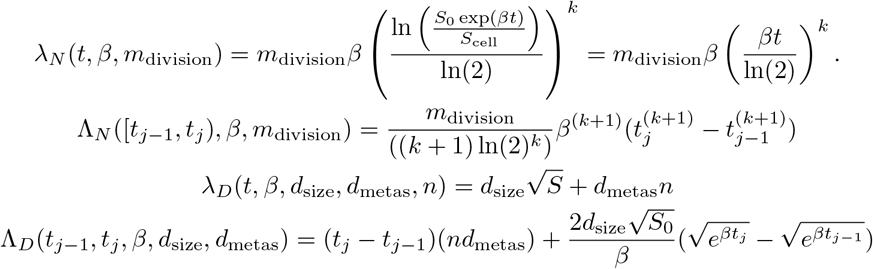

the survival probability over an interval [*t*_*j−*1_, *t*_*j*_) with *N* (*t*_*j−*1_) = *n* takes the following functional forms for a given number of new metastasis *m* ∈ {0, …, 5}.

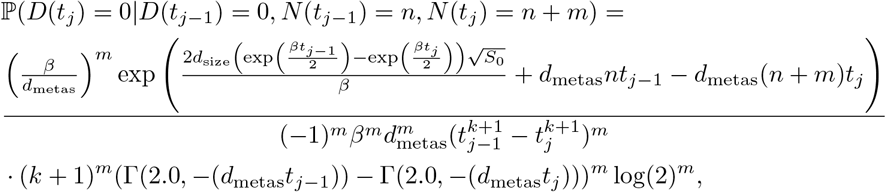

where Γ(*s, x*) denotes the upper incomplete gamma function defined as

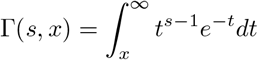

### Exponential Proportional Model

Given the following functions for tumour growth and intensity rates of the Poisson point processes

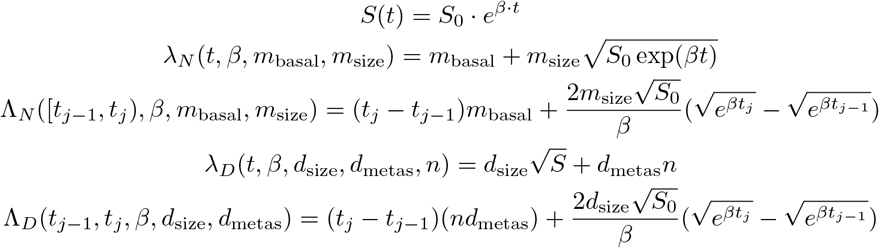

the survival probability over an interval [*t*_*j−*1_, *t*_*j*_) with 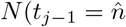 takes the following functional forms for different given number of new metastasis *m* ∈ {0, …, 5}

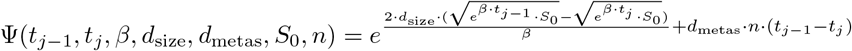

**m=1:**

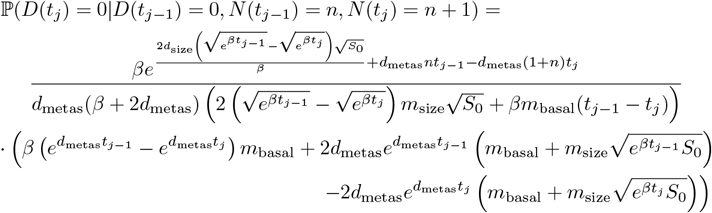

**m=2:**

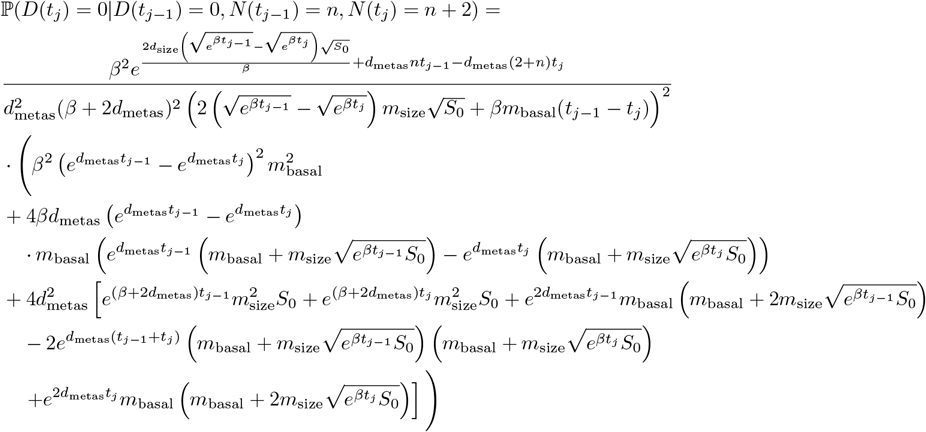

**m=3:**

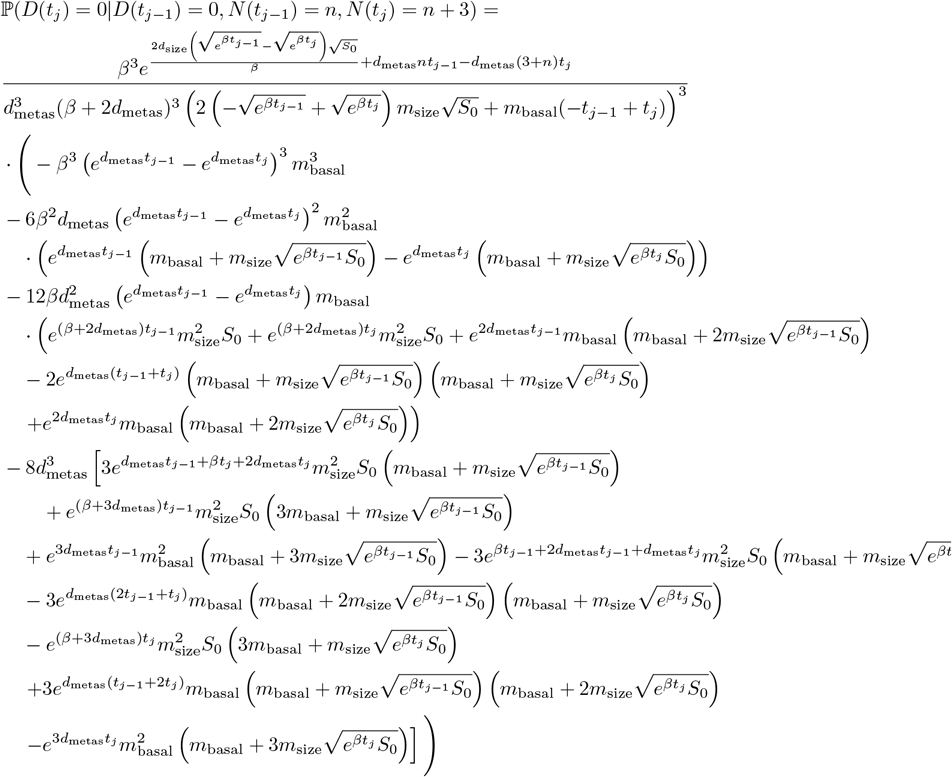

**m=4:**

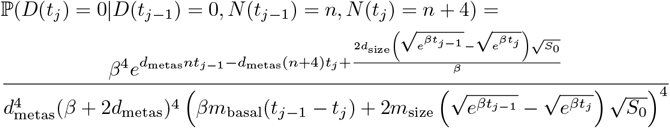

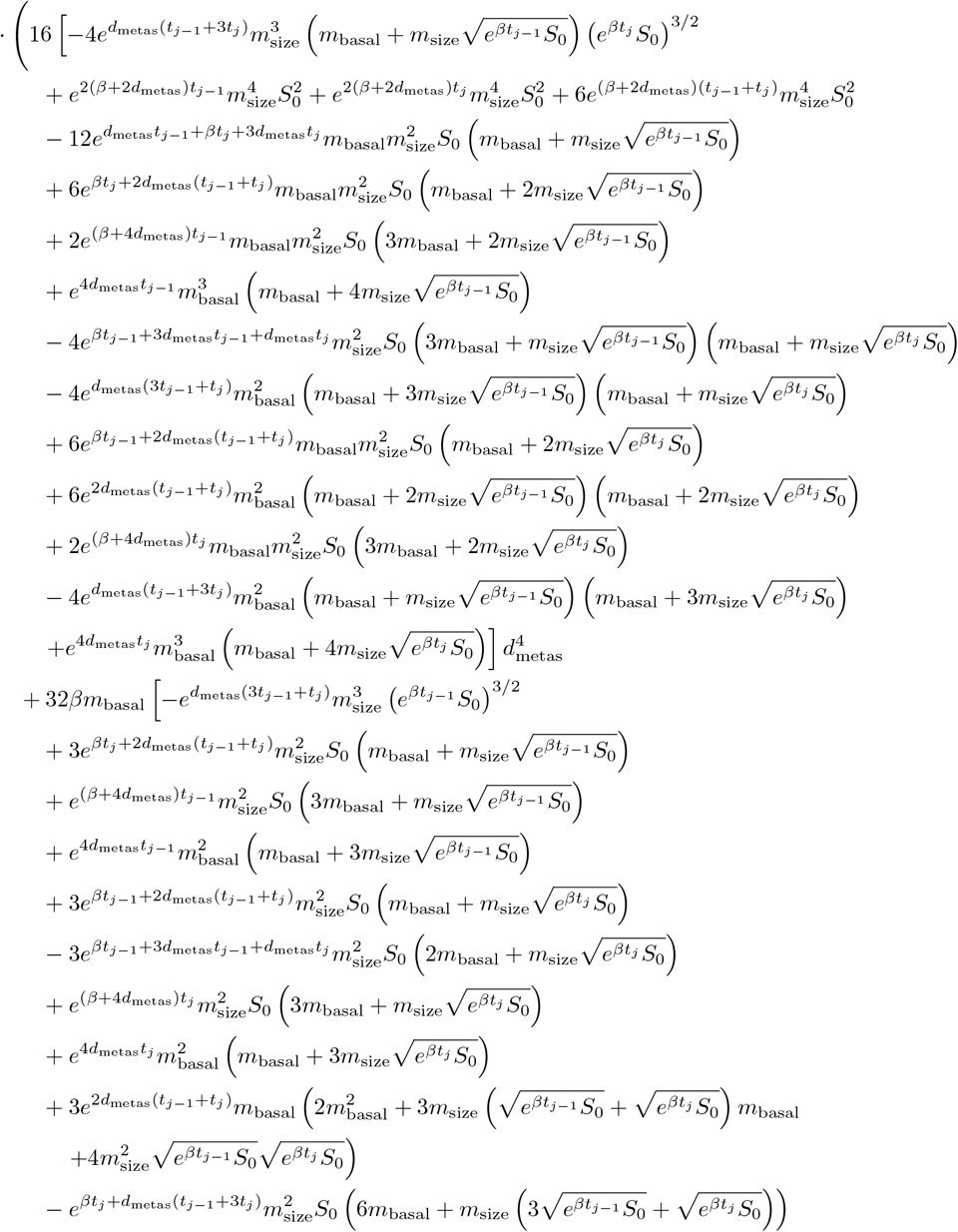

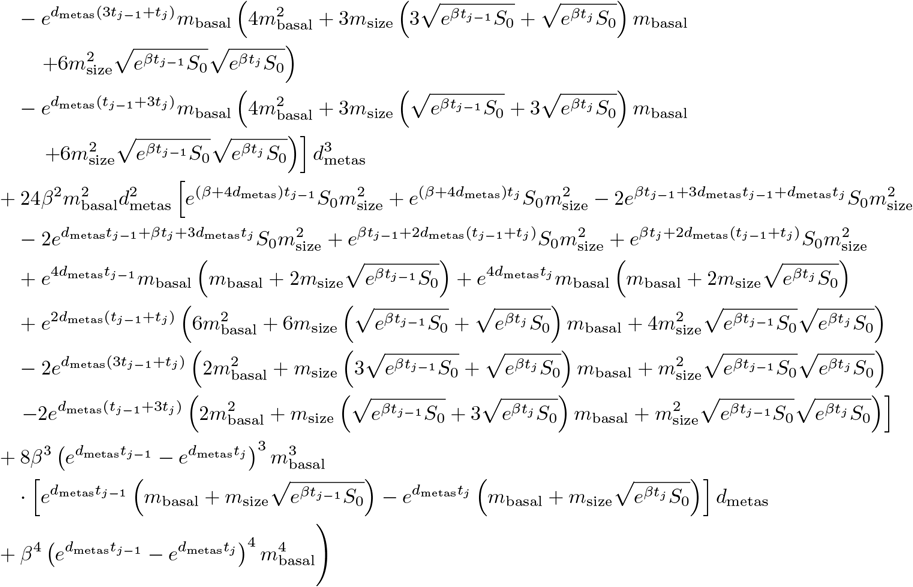

**m=5:**

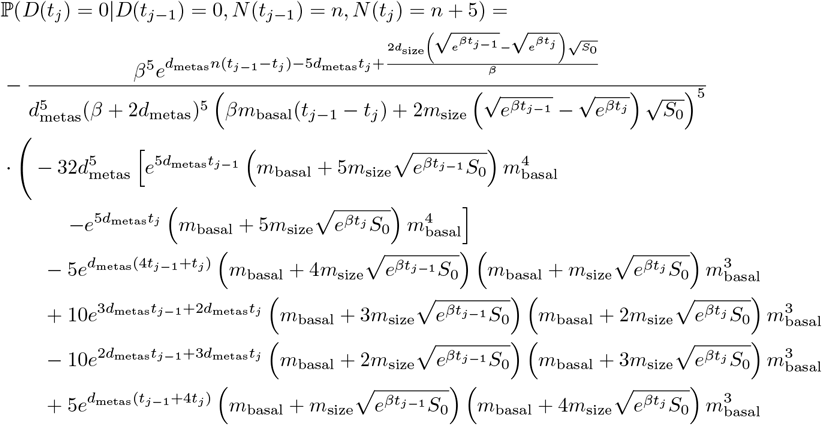

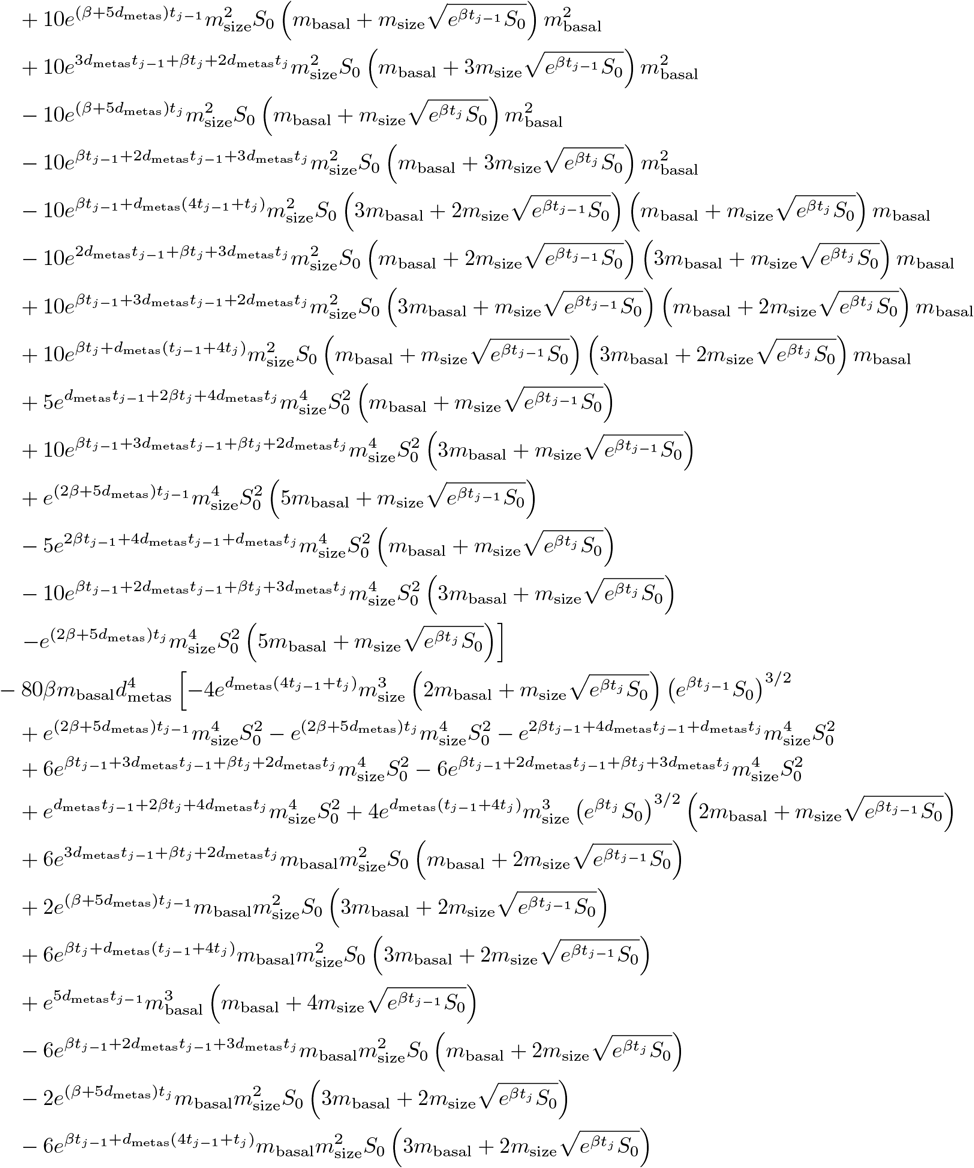

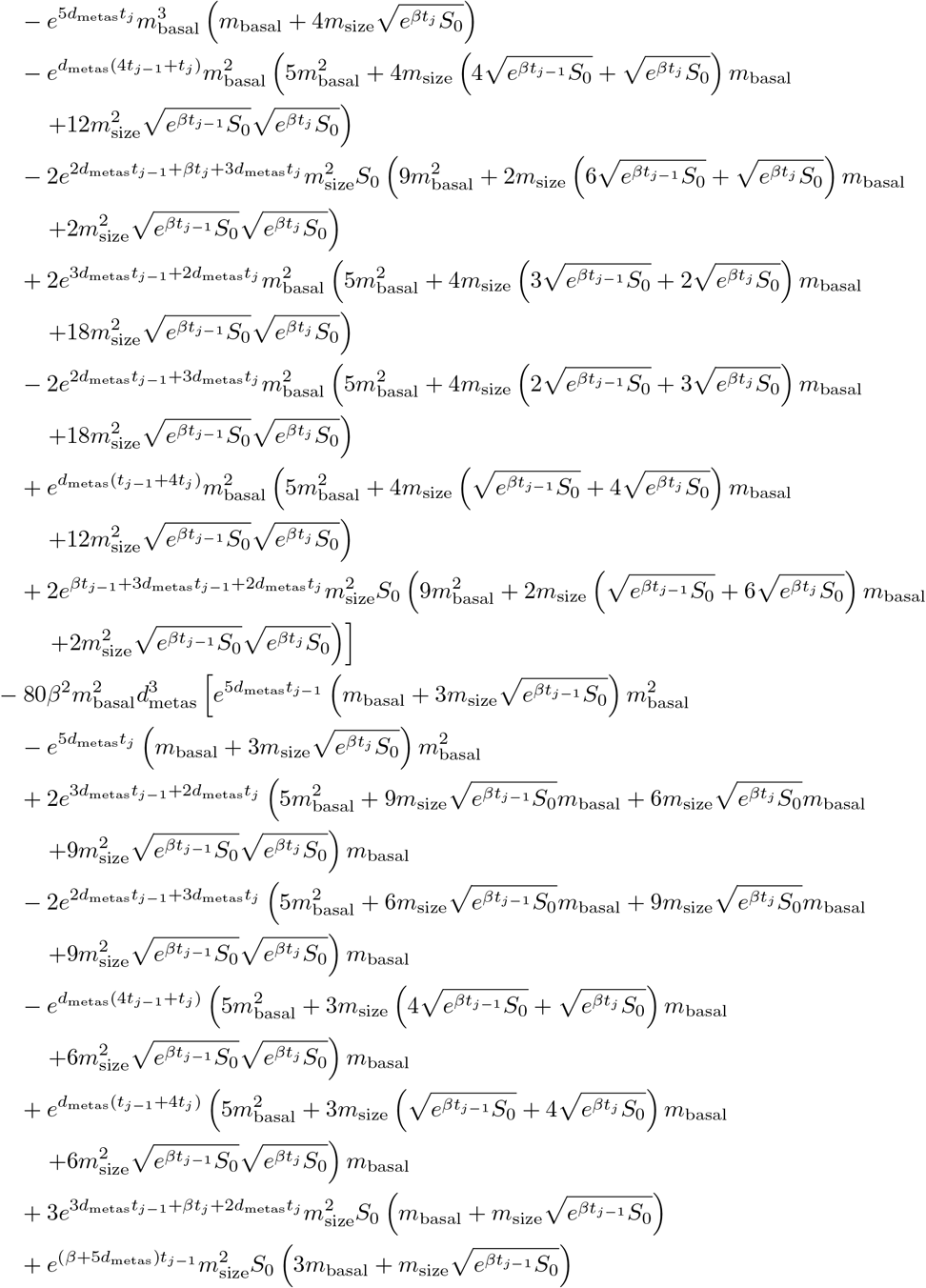

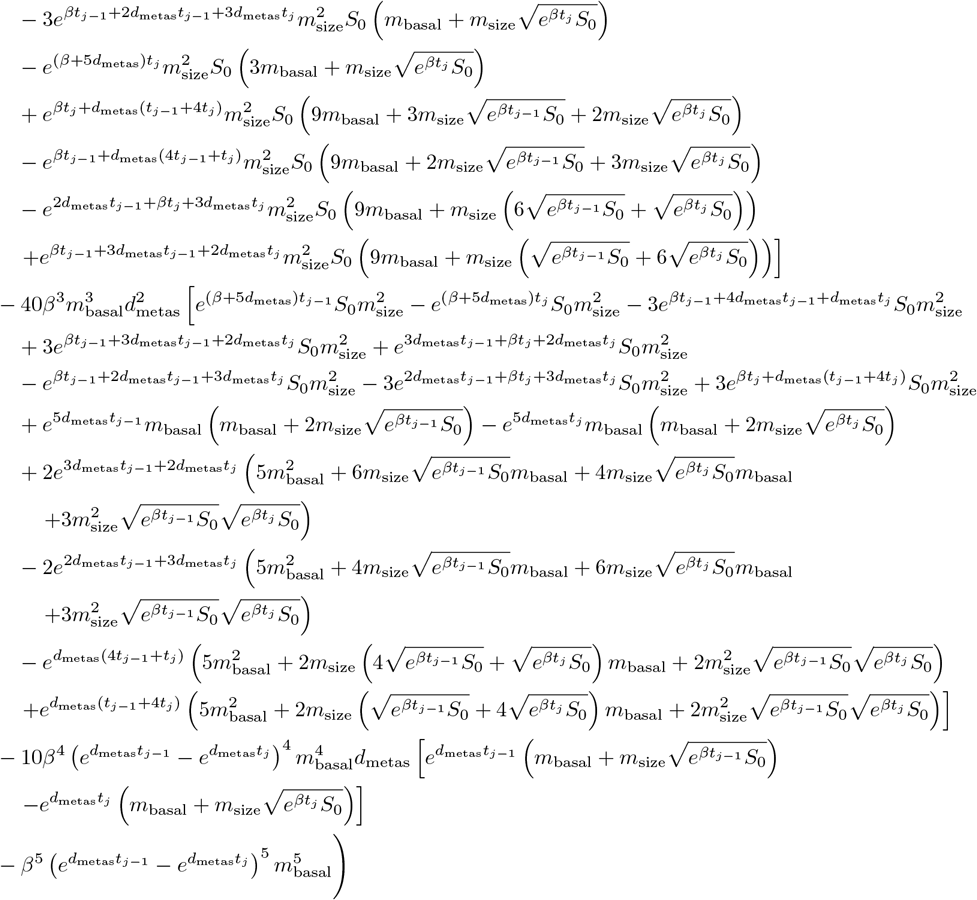

## Footnotes

1 Created in BioRender. Hasenauer, AG. (2024) BioRender.com/k04j643

2 Parametrization given in the Supplementary Information A.2

3 https://www.tumorregister-muenchen.de/en/

4 Compare to eq. 16.13 in (Gabbiani and Cox 2010)

## Notes

https://zenodo.org/records/13839104

https://github.com/vwiela/Combined_Stochastic_Model1.git

### Competing Interest Statement

The authors have declared no competing interest.

### Summary of Updates

The following points were changed in the manuscript: - We streamlined our introduction to better highlight the gaps in the existing model space that are addressed by the proposed work. - Add physiologically structured Gyllenberg-Webb model for the tumour growth process. - Add an analysis of the survival time distribution for the combined model. - Add validation of the proposed modelling framework against data from the Munich Cancer Registry. - Provide likelihood computations for the case of partially or completely missing data. - Enhanced the discussion of the model, its limitations and the outlook onto further research. In addition to these changes in the content, we updated the structure and the section titles as well as the title of our proposed work: - Manuscript title was changed to "A Stochastic Modelling Framework for Cancer Patient Trajectories: Combining Tumour Growth, Metastasis, and Survival" - The Implementation Section was split into the implementation of the model and the implementation of the inference pipeline, where each part was added to the respective sections. - Renamed the "Results" section into "Evaluation of Parameter Inference" to provide clearer guidance for the reader. - Improved the overview figure by adding a part about the mathematical model. - Combined several figures and moved redundant figures into the Supplementary part of the manuscript. These changes are accompanied by additional Supplementary Information and Figures added into the Supplementary part of the manuscript.

